# De novo assembly and annotation of Asiatic lion (*Panthera leo persica*) genome

**DOI:** 10.1101/549790

**Authors:** Siuli Mitra, Ara Sreenivas, Divya Tej Sowpati, Amitha Sampat Kumar, Gowri Awasthi, M. Milner Kumar, Wajeeda Tabusum, Ajay Gaur

## Abstract

We report the first draft of the whole genome assembly of a male Asiatic lion, Atul and whole transcriptomes of five Asiatic lion individuals. Evaluation of genetic diversity placed the Asiatic lion in the lowest bracket of genomic diversity index highlighting the gravity of its conservation status. Comparative analysis with other felids and mammalian genomes unraveled the evolutionary history of Asiatic lion and its position among other felids. The genome is estimated to be 2.3 Gb (Gigabase) long with 62X sequence coverage and is found to have 20,543 protein-coding genes. About 2.66% of the genome is covered by simple sequence repeats (SSRs) and 0.4% is estimated to have segmental duplications. Comparison with seven well annotated genomes indicates the presence of 6,295 single copy orthologs, 4 co-orthologs, 21 paralogs uniquely present in Asiatic lion and 8,024 other orthologs. Assessment of male and female transcriptomes gave a list of genes specifically expressed in the male.

Our genomic analyses provide candidates for phenotypes characteristic to felids and lion, inviting further confirmation of their contribution through population genetic studies. An Asiatic lion-specific expansion is detected in the Cysteine Dioxygenase-I (CDO-I) family that is responsible for taurine biosynthesis in cats. Wilm’s tumor-associated protein (WT1) family, a non-Y chromosome genetic factor underlying male-sex determination and differentiation is found to have undergone expansion, interestingly like that of the human genome. Another protein family, translation machinery-associated protein 7 (TMA7) that has undergone expansion in humans, also expanded in Asiatic lion and can be further investigated as a candidate responsible for mane in lions because of its role in hair follicle morphogenesis.

## INTRODUCTION

The Asiatic lion, *Panthera leo persica*, is one of the most endangered large mammals in the world. A small population of lion in Gir forests in India estimated at 523 (Gujarat Forest Department Census 2015), represents the only known remaining wild population of an animal that historically ranged throughout most of the southeastern and southwestern Asia, extending from Syria to north India, as recently as 200 years ago^1^. The subspecies became extinct in Syria, Iraq, Iran, Afghanistan and Pakistan in the later part of the 19^th^ century. In India too, the lions were once spread across Rajasthan, Gujarat, Haryana, Punjab, Uttar Pradesh, Madhya Pradesh and western Bihar^2,3^. Population pressure and extensive hunting for trophies reduced the Asiatic lions alarmingly less in number^4,5^.

With strengthening conservation measures and habitat recovery by incessant efforts of the Indian state of Gujarat, the number of lions has increased significantly in Gir forests^6^. However, with the entire wild population being confined to a single location, the Asiatic lion is facing an increased risk of extinction because of a very low number and continuous inbreeding^7,8^. Morphological and molecular approaches have been used to study genetics, population structure and evolutionary history of this species^9–11^. The molecular genetic assessment of the Asiatic lions at whole genome level can be utilized for designing further conservation and management strategies for this critically endangered Indian big cat.

Here, we have sequenced, assembled and annotated a draft whole genome of an Asiatic lion male named Atul. We have also characterized the transcriptome of five individual Asiatic lions to obtain data on gene expression and to assist gene annotation.

## RESULTS & DISCUSSION

### Asiatic lion genome sequence and assembly

The genomic DNA was isolated from peripheral blood of a male Asiatic lion named Atul. We used the whole genome sequencing strategy on the HiSeq 2500 platform (Illumina Inc., USA) to reconstruct the genome sequence of Atul (Supplementary Figure L1, Supplementary Tables S1 and S2). The isolated DNA was subjected to sequencing using libraries of varying insert sizes; 150 base pairs (bp), 500bp, 800bp, 6kb (kilo base), 10kb, and 20kb. In total, we generated 231 Gb sequence data with the read lengths of 100 bp and 250 bp for pair-end and mate-pair libraries respectively, achieving a total of 90X fold coverage of the whole genome (Supplementary Table S3). To ensure an accurate assembly, we excluded data from poor libraries, eliminated low-quality reads with erroneous bases or other ambiguous sequence data, and used 152 Gb (62X) high-quality reads for *de novo* assembly of the individual (Table 1, Supplementary Figure L2, L3A and L3B, Supplementary Table S4 and Methods).

**Table 1:** Summary of Asiatic lion genome sequencing and assembly

| SEQUENCING STATISTICS |  |  |  |  |
| --- | --- | --- | --- | --- |
| Insert size |  | Total filtered data (Gb) | Sequence coverage (X) |  |
| 150, 500, 800 bp<br>6, 10, 20 kb |  | 152 | 62 |  |
| ASSEMBLY STATISTICS |  |  |  |  |
|  | > 5kbp | N50 (bp) | Longest stretch | Total size (Gb) |
| Contig | 155,481 | 62, 599 | 1,43,273 | 2.3 |
| Scaffold | 83,167 | 20,864 | 3,83,982 | 2.4 |
| GC content | 39.63% |  |  |  |
| VARIATION STATISTICS |  |  |  |  |
|  | Homozygous | Heterozygous | Total |  |
| Total | 261,257 | 38,517 | 299,774 |  |
| Protein coding genes | 20,543 |  |  |  |
| RNA coding genes | 4,057 |  |  |  |

We assembled the short reads using the de Bruijn graph algorithm of SOAPdenovo (http://soap.genomics.org.cn), a genome assembler used for short read sequences^12^. SOAPdenovo reported a peak sequencing depth at 62X and an estimated genome size of 2.3 Gb by using frequency of 23-mer reads (Supplementary Table S5). The resulting assembly scaffolds have an N50 of 20,864, the longest scaffold being 383 Kb (Supplementary Table S6).

To ensure the quality of assembly, multiple strategies were adopted including mapping lion reads to lion scaffolds, assembling blood transcripts of five Asiatic lion individuals and reference *Feliscatus* 8.0 domestic cat EST sequences (hereafter called Fca-8.0) on to the lion scaffolds which showed a decent coverage of 86% and mapping was quite high at 96.63% (Supplementary Tables S7-S9). Additionally, lion draft genome assembly analyzed by core eukaryotic genes using CEGMA approach revealed that more than 86.8% of conserved genes could be identified in the assembly (Supplementary Table S10), indicating a good coverage and completeness of the core gene repertoire with unique gene sequences in Asiatic lion genome assembly and it is same as that observed in the African lion assembly^13^. The GC content of the Asiatic lion genome was determined to be 39.63% (Supplementary Table S11), close to 41.66% obtained for a domestic cat genome reference genome of Fca-8.0^14^ and 41.4% in Amur tiger^15^ suggesting less bias due to assembly quality.

The draft chromosome of Asiatic lion was constructed by mapping against the domestic cat reference genome of Fca-8.0, the only felid to have a chromosome level genome assembly. The lion genome showed 93.9% similarity to the domestic cat which is similar to findings for tiger (Supplementary Table S12). The divergence times of Felidae radiation estimated by Johnson *et al*.^16^ are suggestive that *P. tigris* diverged from the felid ancestor after *P. leo* and is, hence, expected to show a higher genome similarity^15^ reflected in the slightly lower similarity observed between Asiatic lion and domestic cat genomes (Supplementary Tables S11 and S13-16, Supplementary methods).

### Asiatic lion genome annotation

We compared the pattern of distribution of six important classes of genome repeats in Asiatic lion with four other published felid genomes^13,14,15,17^ and found that the genome coverage for the classes were largely conserved and reflective of the recent divergence except for the small RNAs (Supplementary Figure L4 A and B). The striking difference between small RNAs distribution and a similar distribution among Asiatic lion and cheetah genomes (Supplementary Figure L4 B) might be suggestive of the low evolutionary pressure acting on these groups. Four groups of non-coding RNA could be delineated out of which snRNA (spliceosomal RNA and CD-box) presented the largest genome coverage (1.2%) of a total of 4,057 ncRNA copies (Supplementary Table S17) like the tiger genome (Supplementary Figure L5). We have also searched for the simple sequence repeat (SSR) distribution in lion genome using our exhaustive repeat finding algorithm, PERF^18^ and identified a total of 4.23 million SSRs >=12bp in length. Together, the SSRs cover 2.66% of the Asiatic lion genome. The most frequent SSRs are poly A/T repeats (0.53 million), followed by AG, AAAT, AC, and C repeats. A majority (53.2%) of SSRs are hexamers, whereas, trimers are the least abundant (2.4%) (Supplementary Figure L6A and B).

Annotation of Asiatic lion genome concluded with the prediction of 20,543 protein-coding genes by integrating evidence generated from homology-based, *ab initio* and EST-based approaches (Supplementary Table S18 and S19) compared to 19,043 in Amur leopard^13^, 20,285 in domestic cat^14,^ 20,226 in Amur tiger^15^ and 20,343 in cheetah^17^. A mere 2.63% remained un-annotated (Supplementary Table S20) and the reason can be attributed to the sizable number of annotated pantherine genomes now available for comparison.

### Genetic diversity of Asiatic lion

Felids, in the order Carnivora, have distinctively small population sizes and are also typified by low genetic diversity which holds pertinence in their conservation. Even within felids, the fact, that Asiatic lion lies towards one end of the genomic diversity spectrum, is indicated in a graph showing the rate of heterozygous SNPs per base pair of a genome (Figure 1). This statistics has also been used as the index of genomic diversity in other whole genomes published recently^13,15,19^. Mapping Asiatic lion raw reads to cat genome unraveled 745,184 SNPs and a comparison with genome size gave a rate of 0.000276 heterozygous SNVs per base pair (Supplementary Table S14 and S30). This is lower relative to the other two sub-species of lion, African lion and white lion and comparable with Eurasian lynx which is also facing the risk of extinction^20^. The rate is also much lower than the average (0.00094) reported for Felidae genomic diversity. The low genomic diversity characterized by low rate of heterozygous SNPs per base pair can be attributed to the low effective population size of Asiatic lion in wild^21^ as demonstrated for brown hyena^22^ but a genetic investigation with multiple Asiatic lion individuals will be necessary to confirm this finding.

**Figure 1:**
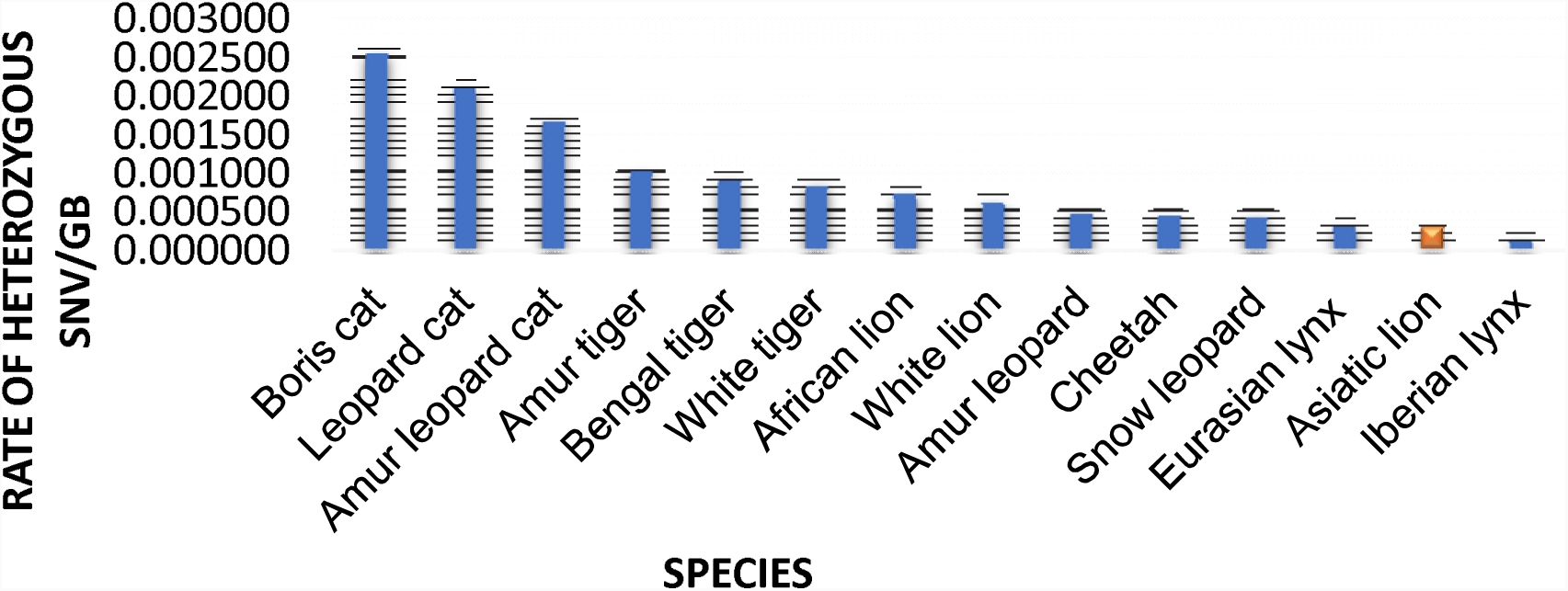
Genomic diversity of Asiatic lion. The rate of heterozygous SNVs per base pair of genome is shown as an index of genomic diversity across felids. The data for other felids has been summarized in Supplementary Table S30.

### Comparative genomics: Asiatic lion perspective

About 0.4% of the lion genome was found to be comprised of segmental duplications. In the synteny analysis by comparison with the domestic cat (*Felis_catus 8.0*) using *nucmer* of MUMmer package^23,24^, a total of 560 blocks of intra-chromosomal and 6,037 blocks of inter-chromosomal rearrangements were found (Supplementary Figure L11, Supplementary Table S32 and S33). A circular map showing syntenic regions between the lion and cat chromosomes is shown in Figure 2.

**Figure 2:**
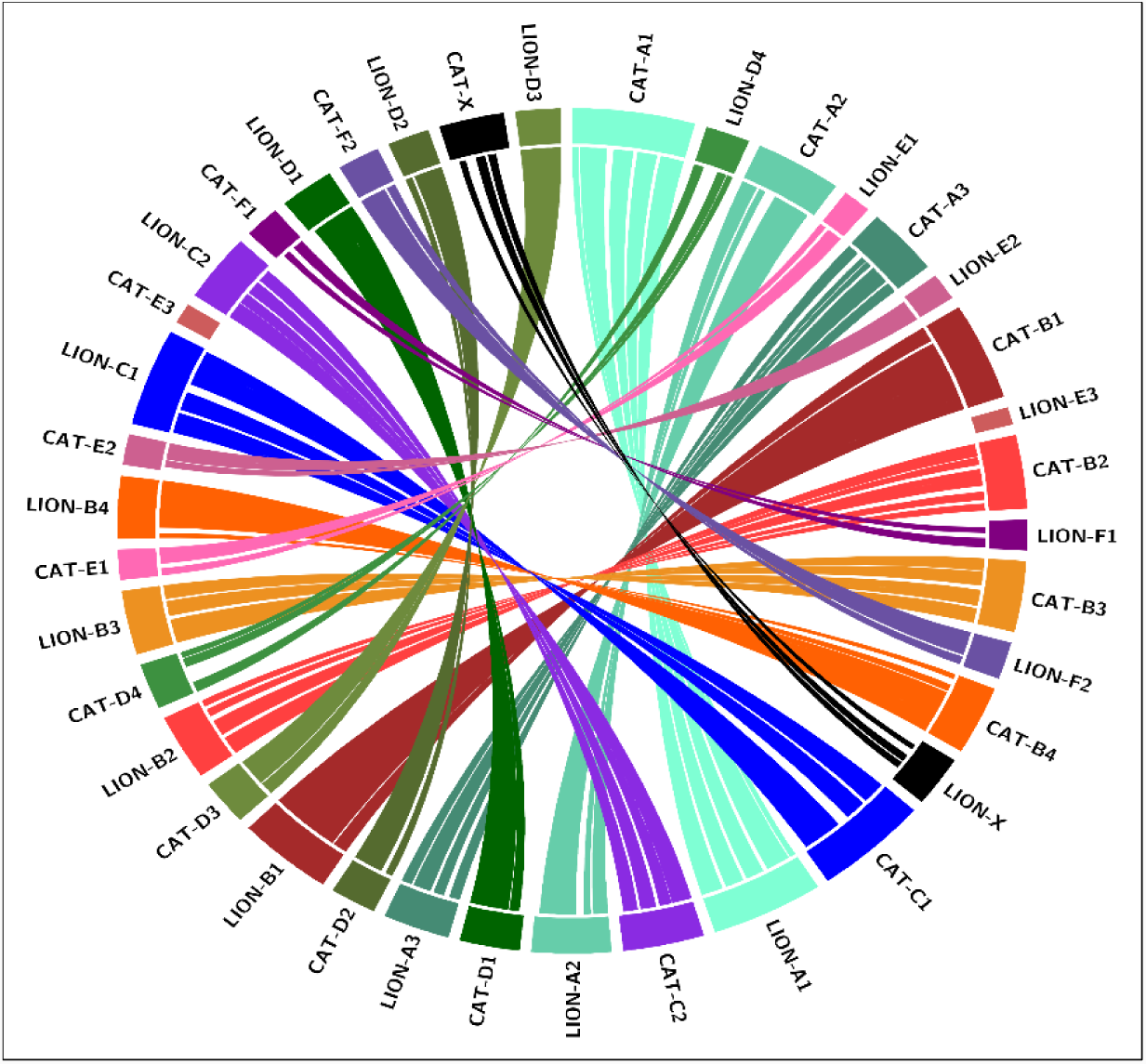
Circos map showing chromosomal synteny between Asiatic lion and domestic cat (*Felis_catus 8.0*) genomes. Each homologous pair of chromosome between the genomes is represented by a distinct color. The gaps in synteny indicate rearrangement.

We identified 20,543 protein-coding genes in the Asiatic lion assembly. The comparison of lion proteome with well annotated available genomes of seven mammals, namely, domestic cat, tiger, dog, panda, human, mouse, and opossum, using OrthoMCL broke up the lion proteome into 6,295 single copy orthologs, 4 co-orthologs, 21 paralogs unique to lion and 8,024 other orthologs (Figure 3 and Supplementary Table S21). Comparison within Felidae revealed 126 protein families, out of which functional annotation for 34 families could be done and most of them were found to be involved in activities related to the metabolic process [GO: 0008152], cellular process [GO:0009987], cell binding [GO: 0005488], cell part [GO: 0044464], catalytic activity andorganelle [GO: 0043226] (Supplementary Table S22). Seventeen genes were identified as lion-specific by Orthofinder, 14 of these were annotated. PANTHER pathway analysis of the gene set showed involvement in signaling pathways namely, JAK/STAT, Interleukin, BMP/activin, Interferon-gamma, TGF-beta, SCW, PDGF, EGF receptor, DPP and DPP-SCW (Supplementary Table S23). Lineage-specific counts of protein families expanding, contracting and propensity for gains and losses were enumerated using the Wagner parsimony algorithm of COUNT software^25,26^. An asymmetric version of the Wagner parsimony used penalized genomic gain and losses differentially. The phylogenetic tree obtained from PhyML and a multiple sequence alignment file were input for the analyses. The gain penalty was set at a strict value of 2.5. The protein families found to be expanded in Asiatic lion were functionally classified using Gene Ontology, KEGG, and PANTHER (Supplementary Table S24, S25 and S27) and included G protein-coupled receptors expansion of which has typically been reported among other felids^27^. A total of 240 lion protein-coding genes were determined as positively selected and were classified according to their biological, cellular and molecular functions (Supplementary Figure L10). Further pathway analysis was carried out using PANTHER (Supplementary Table S28).

**Figure 3:**
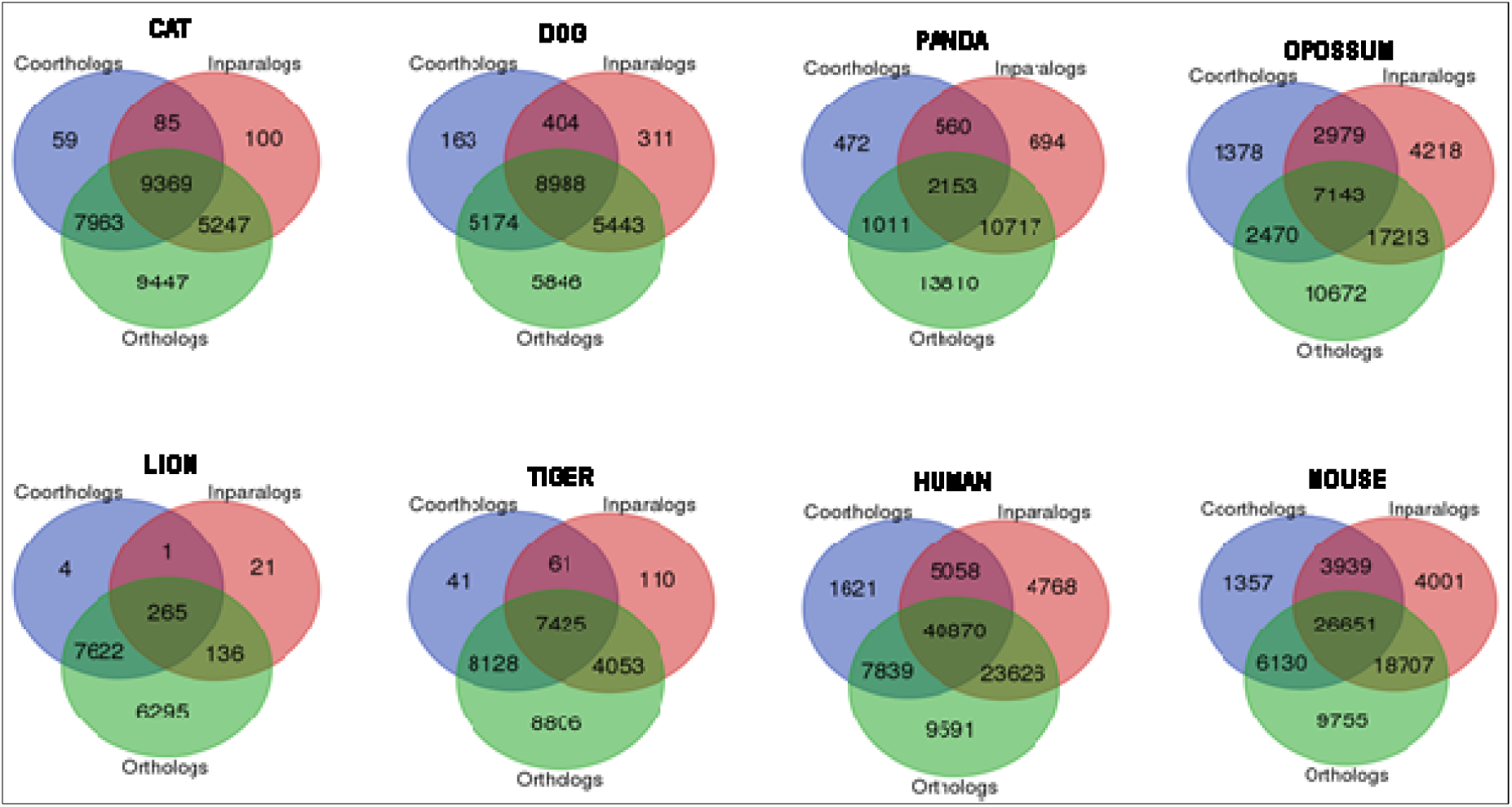
Venn diagram showing orthologous protein families for eight species. Orthologs and co-orthologs are result of speciation events and in-paralogs the result of a duplication event within a genome.

The rate of heterozygous SNVs per base pair of the genome was calculated for Asiatic lion and compared with 11 other whole genomes to decipher the genomic diversity of the former among felids (Supplementary Table S29).

#### Evolution in specific protein families and genes

Inferences drawn based on protein families expanded and contracted in a species proteome, might be reflective of the specific adaptations the organism (or a lineage) has undergone and is largely because of gene duplication, divergence or recombination^28^. The asymmetric application of Wagner’s parsimony algorithm was used to delineate the protein-families that underwent these lineage-specific evolutionary changes in Asiatic lion. Twenty-four protein families expanded while 792 underwent contraction in this species after its divergence from the common ancestor shared with the tiger lineage (Supplementary Table S26). Phylogenomic analysis showed the origin of a lion to be a later event as compared to other felids at 10.5 Mya (95% CI: 10.18-10.92Mya) (Figure 4).

**Figure 4:**
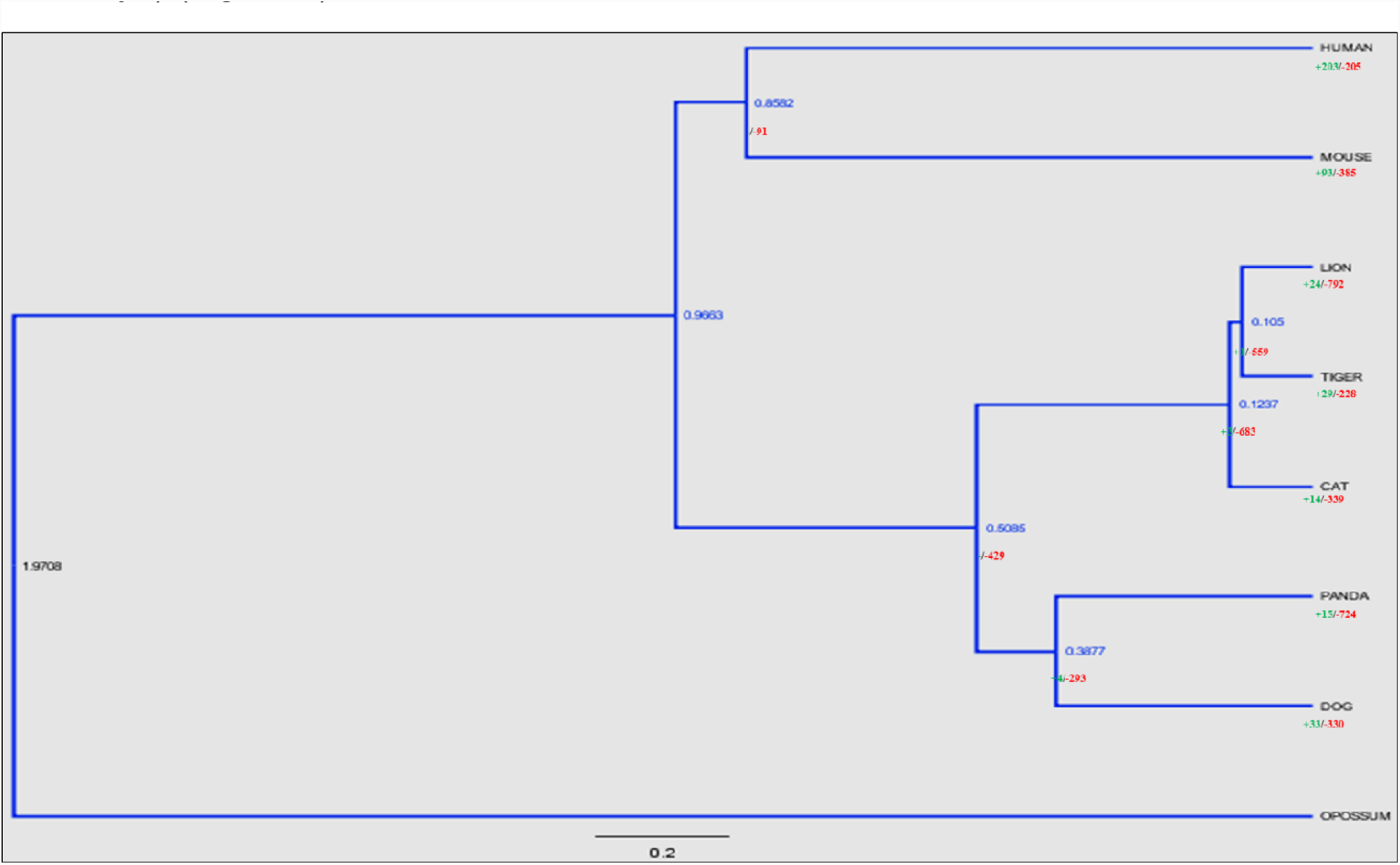
Divergence time estimates and protein families’expansion (+) and contraction (-) in seven mammalian proteomes including Asiatic lion.

The Cysteine Dioxygenase-I (CDO-I) family (PTHR12918) found to be expanded in the Asiatic lion (Supplementary Table S26), is involved in the biosynthesis of the amino acid taurine which is known to be an essential nutrient in a cat’s diet and its absence causes retinal degeneration and blindness^29^. Along with cysteine sulfuric acid decarboxylase (CDAS), CDO-I is responsible in the formation of taurine from cysteine while the former is the rate-limiting enzyme in this process. Thus, an expansion specific to Asiatic lion can only be explained by a genomic gain that does not lead to a change in the CDO-I enzyme activity to enhance taurine biosynthesis.

The Wilm’s tumor-associated protein (WT1) family is a known non-Y chromosome genetic factor responsible for male-sex determination and differentiation in animals^30^ and mutations in this gene are known to cause urogenital abnormalities in humans^31^. The family was also demonstrated to have undergone a genomic gain in human genome and absence of the same in the closely related tiger in our analysis. The WT1 family was found to have undergone expansion in Asiatic lion after its divergence from the pantherine ancestor although a gender-specific difference in gene expression was not observed in our analysis. Knock-out analyses in mice have shown reduced gene expression of male-specific markers^32^.

The translation machinery-associated protein 7 (TMA7) underwent expansion in Asiatic lion. It was interestingly found to have undergone gain in human while experiencing a loss in the more closely related domestic cat and Amur tiger (Supplementary Table S26). Null yeast strains with TMA7 gene deletions have been shown to have altered rates of protein synthesis^33^. HSPC016, a TMA7 clone, was shown to be differentially expressed in dermal papilla cells (DPC). The DPC are pivotal in hair follicle generation and morphogenesis. The inhibition of HSPC016 led to slower growth and non-aggregative behavior of DPC^34^. The linkage between TMA7 and hair follicle morphogenesis, its expansion in Asiatic lion and human genomes both of which are known to have male facial hair as a common feature makes it an important candidate for examination of a possible role in mane formation of lions.

The comparative analysis helped to provide genomic candidates to explain phenotypic specialization in felids. For instance, the pathway analysis of protein families expanded in Asiatic lion showed the participation of ionotropic glutamate receptor and metabotropic glutamate receptor group III (PTHR18966:SF269) pathways. Glutamate is one of the neurotransmitters that play a key role in synaptic transmission in the vertebrate retina^35^. Its role in low-light vision of cat retina has been specifically investigated^36^ which is a sensory adaptation characteristic of felines as it assists their need to locate the prey in low-light conditions^37^. The SLC6A16 belonging to the orphan sodium and chloride dependent neurotransmitter transporter family, NTT5, was found to be positively selected in the lion genome. Positive selection in this gene might be contributory to meet the necessity of sharper vision in lion as also among felids, in general.

Gender-biased gene expression^38^ has been observed in transcriptome studies across distinct species and the findings have been instrumental in making inferences on important sex-specific traits. The analysis of differential gene expression between three male and two female Asiatic lion individuals revealed four protein coding genes and one region coding for a protein domain of ATP dependent RNA helicase DDX3Y-like protein, all except, one Y-linked, to be enriched among males (Post FC> 2). The other male-enriched genes are Y-linked ubiquitin-specific protease 9 (USP9Y), eukaryotic translation initiation factors 2, subunit 3 (EIF2S3Y) and lysine demethylase 5D (KDM5D) and X-linked Zinc finger protein ZFX.

We have presented the first ever whole genome sequence of Asiatic lion. Findings on genome organization and evolutionary divergence analyses are concordant with those obtained for felid genomes in earlier complete genome studies. Low genomic variation found in Asiatic lion here is typically reported for species with low population sizes and at the brink of extinction which is important from the conservation point of view. Genomic and gene evolution findings in the study especially, relevant to features characteristic of lion and felids, need further corroboration through studies on multiple individuals before they can be used for conservation and management of Asiatic lion.

## METHODS

### Sequencing and *de novo* assembly of the Asiatic lion (*Panthera leo persica)* genome

Genomic DNA was isolated from a fresh blood sample of a male Asiatic lion, Atul housed in Nehru Zoological Park, Hyderabad, India using NucleoSpin^®^ Blood L (Machery-Nagel, GmbH & Co., Germany) followed by a quality and quantity check to ensure high molecular weight DNA for library preparation. Libraries with short insert paired-end reads (150bp, 500bp and 800bp) and long insert mate-pair reads (4-6Kb, 8-10Kb and 1-20Kb) were sequenced using HiSeq 2500 (Illumina Inc., USA) at SciGenom Labs, Cochin, India.

The raw sequencing data obtained was refined by Cutadapt^39^ and mate-pair data extracted using NxTrim^40^ prior to assembly. The genomic contigs were assembled by SOAPdenovo^41^ and CLC Genomics Workbench 9.5.3 (CLC bio, Aarhus, Denmark) and scaffolded using CEGMApipeline^42^. Scaffolds obtained from SSPACE^43^ for CLC assembly were chosen for downstream analysis. A de Bruijn graph constructed with 23-mer frequency of read data gave an estimated genome size of 2.3 Gb (Supplementary Table S5; Supplementary Figure L2). To check the quality of assembly, the weighted mean of assembled contigs (N50) was determined (Supplementary Table S12, Supplementary Figure L3) and Asiatic lion transcripts were mapped to cat ESTs (Supplementary Table S15).

### Genome annotation and Gene evolution

The Asiatic lion genome was scanned for tandem repeats using Tandem Repeats Finder version 4.09^44^. Homology-based approach was applied supplemented with information available in Repbase version 21.06^45^ to identify transposable elements (TEs). The database was used to locate repeats using Repeat Masker version 4.0.6. A SINE-masked genome was scanned for tRNA using tRNAscan-SE version 2.0^46^ for determining reliable tRNA positions. snRNA and miRNA were discovered by a BLAST alignment followed by running INFERNAL^47^ to search for putative sequences in the Rfam database Release 12.1. Simple Sequence Repeats >=12bp in length were identified using PERF^18^.

The Asiatic lion genome annotation was carried out by following an in-house pipeline CANoPI (Contig Annotator Pipeline) maintain by Scigenom Ltd., for comparison with UniProt using BLASTP, organism annotation, gene and protein annotation to the matched CDSs, gene ontology and pathway annotation.

Phylogenomic analyses were shown to delineate the position of Asiatic lion in the mammalian and felid phylogenies. To accomplish this, we used published whole genomes of human (GRCh38.p9), mouse (GRCm38.p5), cheetah (*AciJub1.0*), tiger (*PanTig 1.0*), domestic cat (*Felis_catus-8.0*), panda (*aiMel1.0*), dog (*CanFam3.1*) and opossum (*monDom5.0*) was reconstructed using concatenated amino acid sequences. The phylogenomics was reconstructed using BioNJ^48^ of PhyML^49^ followed by fastNNI^50^ for tree optimization and LG model of substitution^51^ was applied. TREEVIEW was used for the visualization of resulting trees. (Supplementary Figure L8 and L9). Divergence dating was done using MCMCTREE implemented in PAML version 4.0^52^.

The proteome of Asiatic lion was again compared with the well-annotated proteomes of seven mammals, domestic cat, tiger, dog, panda, human, mouse, and opossum to delineate the extent of gene orthology among these organisms using TreeFam^53^ and OrthoMCL^54^ methodologies to predict orthologs, co-orthologs and paralogs. The Pfam^55^ was used to predict the number of protein families (Supplementary Table: S21). These analyses helped to identify gene function in Asiatic lion genome by scrutinizing conserved regions among gene orthologs. Since the organisms being examined belong to different orders (*Carnivora, Rodentia, Primata, Didelphimorphia*), the analysis was helpful in studying the extent of protein evolution in them.

The ratio of rates of non-synonymous (dN) and synonymous substitutions (dS), ω was computed using the CODEML program of the PAML package^56^ from the alignment of annotated genomes of Asiatic lion, tiger, cat, dog, mouse, panda and opossum obtained by PRANK^57,58^. Only 1:1 orthologs were used for this analysis. The branch-site model in which ω can vary at specific sites of a gene sequence and branches of phylogeny was applied^59^.

The blood transcriptomes of three male Asiatic lions and two female Asiatic lions were aligned using STAR, which uses a sequential maximum mappable seed search followed by seed clustering and stitching procedure^60^. RNA-Seq by Expectation Maximization (RSEM) was applied to evaluate and carry out a differential analysis^61^ allowing a false discovery rate less than 0.05. The number of paired reads (expected counts) was estimated for genes exhibiting a 2-fold change were considered as differentially expressed in male samples (Supplementary Table - S30). Male-specific genes were selected based on a minimum value of 2 in PostFC which is the posterior fold change for a gene/transcript each gene.

### Draft chromosomes of Asiatic lion

The Asiatic lion genome contigs were aligned with the genome of domestic cat (*Felis_catus 8.0*) as reference using bwa-mem program of BWA^62^, to determine the chromosome location and order of scaffolds. EMBOSS^63^ was used to merge two overlapping scaffolds. Chromosomes and pseudo-molecules were reconstructed based on the synteny observed between the Asiatic lion and domestic cat genomes.

### Chromosomal synteny

The Whole Genome Assembly Comparison (WGAC) was done to find segmental duplications in the Asiatic lion genome. The synteny between Asiatic lion and domestic cat chromosomes was reconstructed by assembling the genomes using *nucmer* of MUMmer package^64^. A region/block of Asiatic lion chromosome mapped distant to domestic cat chromosomal locations was noted as an inter- or intra-chromosomal rearrangement event.

## Data Availability

Raw DNA and RNA sequencing data and genome assembly are available at NCBI Sequence Read Archive database (SRP103576) under the Bioproject accession number PRJNA379375 and PRJNA384102.

## Declarations

The authors declare no conflict of interests.

## Acknowledgements

We acknowledge the incessant support of the Dr. Rakesh Mishra, Director, CCMB, Dr. Ch. Mohan Rao, Former Director, CCMB, Dr. Karthikeyan Vasudevan, Scientist In-Charge, LaCONES and Dr. S Shivaji, Former Scientist In-Charge, LaCONES. We thank the PCCF (WL) & CWLW, Telangana Forest Department for necessary permissions for sample collection. We also thank Director, Curator, Veterinary Doctors and staff of Nehru Zoological Park, Hyderabad, India, for coordinating and providing all support during sample collection. We thank our colleagues Dr. Sambasiva and Dr. Sadanand Sontakke for collecting blood samples. We sincerely acknowledge Dr. Rakesh Mishra, Dr. K. Thangaraj, Dr. Jyotsna Dhawan, Dr. D. P. Kasbekar and Dr. Swasti Raychaudhuri for their critical comments and valuable suggestions which have helped to improve the manuscript significantly. Ms. Vaishnavi Kunteepuram, Ms. Drashti Parmar and Mr. Noopur Modi are also acknowledged for their invaluable help during sample collection and data analysis. We gratefully acknowledge CSIR, India for funding (Grant No. BSC0207).

