## Supplemental Methods for "De novo assembly and annotation of Asiatic lion (*Panthera leo persica*) genome"

**Preparation of sample for sequencing:**

The blood sample of an Asiatic lion individual named Atul was obtained from Nehru Zoological Park, Hyderabad (vide permission letter from Chief Wildlife Warden and Special Principal Chief Conservator of Forests-Dev., Telangana, letter Rc. No. 43505/2013/WL-1, dated 12/11/2014, (**Supplementary Table S1)**. DNA was isolated using the lysis method of NucleoSpin® Blood L (Machery-Nagel, GmbH & Co., Germany) following the manufacturer's recommendations. The quality and quantity of the sample was checked using NanoDrop ND 1000 (ThermoFisher Scientific, USA), Qubit (ThermoFisher Scientific, USA) and gel electrophoresis **(Supplementary Table S2 and Figure L1)** to make sure that the DNA obtained is sufficient and of superior quality for library preparation.

**Asiatic lion genome sequencing:**

Libraries of different insert sizes were constructed at SciGenom Labs, Cochin, India. Three short insert paired-end libraries (150bp, 500bp and 800bp) and three long insert mate-pair libraries (4-6Kb, 8-10Kb, and 1-20Kb) were sequenced using HiSeq 2500 (Illumina Inc., USA) **(Supplementary Table S3)**. A total of 231 Gb (Gigabase) data was generated as a result.

**Raw read filtering for *de novo* assembly:**

The post-sequencing analysis was earmarked with filtering steps for data quality control and comprised removal of low quality reads, trimming of adapter sequences and sequencing error correction to avoid a low-quality genome assembly. After these steps, the six libraries carrying varied insert sizes remained with that of 152 Gb data (**Supplementary Table S4)**.

***De novo* assembly of the Asiatic lion genome:**

The *de novo* assembly of the filtered reads was carried out using the De Bruijn graph-based approach in the SOAPdenovo software[42] (http://soap.genomics.org.cn/soapdenovo.html). The contigs were further organized into scaffolds. Scaffolding was done using SOAP scaffold and SSPACE[44] with the help of the domestic cat genome owing to its phylogenetic proximity to the Asiatic lion and being the only felid with a chromosome level assembly.A total of 2.46 GB of scaffolds were assembled.

The genome size of Asiatic lion was estimated by using varied K-mer sizes of 17bp, 21bp and 23bp (**Supplementary Tables S5 and S6)**. The estimation was carried out using BBMap (https://sourceforge.net/projects/bbmap/).

**Asiatic Lion blood Transcriptome:**

The quality of the Asiatic lion genome assembly was evaluated by sequencing the transcriptome of five Asiatic lion individuals: Ajay, Atul, Rita, Sonia and Viswas using HiSeq2000 (Illumina Inc. USA). The Trinity algorithm[65] (http://trinityrnaseq.sourceforge.net/) was applied to assemble the Asiatic lion transcriptome as the latter is specifically used for *de novo* assembly of transcripts **(Supplementary Table S7)**. Trinity was chosen as it recovers full-length transcripts with efficiency that is equal across varied expression levels. It depends on the De Bruijn graphs to reconstruct the transcriptome by recovering alternate spliced isoform and recently duplicated genes.

Following that, Blat was used to align the assembled transcripts with the Asiatic lion assembly. The proportion of assembled transcripts covered in the Asiatic lion scaffolds was about 81% **(Supplementary Table S8)**.

Another step taken to examine the quality of the assembly, sequences of domestic cat EST were recovered from (http://hgdownload.cse.ucsc.edu/goldenPath/felCat4/bigZips/est.fa.gz) to be mapped to the Asiatic lion genome sequence. About 86% of the domestic cat ESTs was covered by the lion assembly while the mapped percent was quite high at 96.63% (**Supplementary Table S9**). Additionally, Asiatic lion draft genome assembly analyzed by core eukaryotic genes using CEGMA[43] approach revealed that >86.8% of conserved genes in the assembly **(Supplementary Table S10)** indicating that good coverage and completeness with unique gene sequences in Asiatic lion genome assembly and it is same as that observed in the African lion assembly. The GC content of the Asiatic lion genome was determined to be 39.63% close to 41.66% obtained for a domestic cat and 41.4% in Amur tiger suggesting less bias due to assembly quality **(Supplementary Table S11).**

**Draft Chromosomes of Asiatic lion:**

The draft chromosomes of Asiatic lion were constructed by mapping it to the reference genome of domestic cat (*Felis_catus* 8.0). The phylogenetic proximity of the two species is suggestive of the fact that the chromosomal location and ordering information of the lion scaffolds could be derived from the domestic cat genome **(Supplementary Table S12)**.

Homozygous variations (88% of SNVs and 11% of indels), found by mapping the Asiatic lion short reads to Asiatic lion scaffolds, were replaced with the nucleotides of the cleaned reads **(Supplementary Tables S13 and S14)**. We aligned the short reads of Asiatic lion to the scaffolds using Bowtie2-2.2.5[66] with very sensitive option, and variations using the SAMtools-1.2[67]. Further, when the Asiatic lion raw reads were mapped to the Asiatic lion scaffold 96.65% of the reads mapped to the lion assembly generated earlier. Also, 221,397 homozygous SNVs (73.85%) and 22,790 heterozygous SNVs (7.60%) were detected.

The corrected Asiatic lion scaffolds were mapped to the domestic cat genome (*Felis_catus-8.0*), using Burrows Wheeler Aligner, BWA[68], to determine the chromosomal location and order of the scaffolds. Two overlapping scaffolds without any mismatch were merged using megamerger of EMBOSS package[69] and non-overlapping scaffolds were joined using 100 'X' characters. In addition, the Asiatic lion short reads were aligned to the domestic cat genome using BWA with default options, and continuous mapped regions (consensus sequences) were extracted. A total of 41,532,416 homozygous SNVs were used to generate the consensus sequences **(Supplementary Table 15)**. Finally, 122,195 of 134, 284 scaffolds (91% of the total scaffold length), were placed in the Asiatic lion chromosomes. **(Supplementary Tables S15 and S16)**.

**Assembly quality:**

The N50 is the most appropriate parameter to adjudge the contiguity of an assembly and is the weighted mean of assembled contigs. The lion genome was assembled and contigs were constructed using CLC Genomics Workbench 9.5.3 (CLC bio, Aarhus, Denmark,https://www.qiagenbioinformatics.com/), which utilizes the SIMD instructions to parallelize and accelerate the assembly algorithms, making the program the fastest next generation sequencing assembler at present. For scaffolding of the contigs, SSPACE was used and using this software, a total of 2.46 Gb of scaffolds were assembled. The N50 of the scaffolds was 36,647bp, and the size of the longest scaffold was 383,982 bp **(Supplementary Table S6)**.

**Repeat annotation:**

Repeated sequences are patterns of nucleic acids (DNA or RNA) that occur in multiple copies throughout the genome. They fall into three broad categories: (a) Terminal (Simple) repeats, (b) Tandem repeats and, (c) Interspersed repeats. Here, we have analyzed the microsatellite distribution in Asiatic lion using a comprehensive SSR mining tool, PERF[18] and also Tandem Repeat Finder (TRF) version 4.09[44] was applied to find tandem repeats present in the scaffolds, and other repeats were also identified using homology-based approaches with Repbase version 21.06[70] which is a repository for known repeats. RepeatMasker version 4.0.6 ( http://www.repeatmasker.org) was applied for DNA-level identification in combination with Repbase **(Supplementary Figures L4 A and B, L6 A and B).**

**Gene prediction:**

The prediction of Asiatic lion genes is described in four steps:

a) ***De novo* prediction:** *de novo* prediction was performed on the repeat masked genome of Asiatic lion using AUGUSTUS version 3.1.0[71] which is based on Hidden Markov Model (HMM). To perform the gene prediction, the training set of lion was created by using cat proteome and the lion assembly due to the unavailability of a trained data set. Scipio constructed gene structure from assembly of lion and proteins of *Felis catus* from UniProt[72].

Those gene structures were used for the creation of training set. A hint file was also generated from the transcriptome data of lion and incorporated in to the gene prediction.

b) **Homology-based prediction**: Homologous proteins of other species (cat, tiger, dog, human and mouse from NCBI) were mapped to the lion scaffolds using tBlastn (Blast 2.2.31) with an e-value cut offequal to 1E-3. The aligned scaffolds as well as its query proteins were then filtered and passed to Exonerate version 2.2.0[73] to search for accurate spliced alignments.

c) **EST-based prediction:** ESTs of *Felis catus* were taken and aligned to the genome of Asiatic lion using BLAT-35x1 (identity ≥ 0.90, coverage ≥ 0.90)[74] to generate spliced alignments.

d) **Integration evidence:** Source evidences generated from the three approaches mentioned above was integrated using EvidenceModeler-1.1.1[75] to produce consensus gene set. The summary evidences were created by performing blastn with all the predicted sequences from *ab initio*, homology and ESTs against EVM predicted CDS **(Supplementary Tables S18 and 19)**.

**Gene function annotation:**

Gene functions were assigned according to the best alignment match with SwissProt and TrEMBL Mammalian database from UniProt[76]. The annotation has been carried out using an in-house pipeline (CANoPI – Contig Annotator Pipeline). Briefly, we performed the following steps for annotation **(Supplementary Table 20)**.

- Comparison with UniProt Mammalian database using BLASTp program
- Organism annotation
- Gene and protein annotation to the matched CDSs
- Gene ontology annotation
- Pathway annotation

**Non-coding RNAs:**

We detected four types of non-coding RNA in the Lion genome by searching databases using the whole genome sequence, tRNAscan-SE version 2.0[46] was performed on a SINE pre-masked genome to search for reliable tRNA positions. snRNA and miRNA were sought through a two-step method. After alignment with BLAST[77], INFERNAL 1.1.1 was used to search for putative sequences in the Rfam database release 12.1[78] **(Supplementary Table S17)**.

**GENE EVOLUTION:**

**Phylogenomic analysis:**

**a) Asiatic lion and genomes of other published mammalian genomes**

A species tree depicting the inter-relationship of Asiatic lion and other mammalian genomes human (GRCh38.p9), mouse (GRCm38.p5), cheetah (aciJub1.0), tiger (PanTig 1.0), cat (Felis_catus-8.0), panda (aiMel1.0), dog (CanFam3.1) and opossum (monDom5.0) was reconstructed using concatenated amino acid sequences in the PHYLIP format using PhyML which applies BioNJ[49] which is a fast distance based method and helps in computing the initial tree quickly. Following that, fastNNI (nearest neighbor interchange) operations are applied for tree optimization. The alternative NNI trees are evaluated and arranged according to their ML value. NNIs with maximum ML values are applied to the current tree and fastNNI continued till there is no scope for ML improvement. The LG substitution model was used. The resulting phylogenetic tree was visualized using TREEVIEW[79].

**b) Asiatic lion with published genomes of other felids**

The phylogenetic relationships of felids were reconstructed using four published genomes of felids (Asiatic lion, tiger, cat and cheetah) and panda and dog as outgroups. The tree was again reconstructed using the LG model of substitution in PhyML.

**c) Divergence time estimation and rates of substitution**

The divergence time for the eight mammalian genomes including Asiatic lion was estimated using amino acid sequences in the MCMCTREEimplemented in the PAML package v 4.0[52] . The calibration time used for the human-mouse divergence (90Mya) was taken from the TimeTreedatabase (http://www.timetree.org). The time of divergence between the Asiatic lion and Amur tiger was determined to be 10.5Mya (95% CI: 10.18-10.92Mya).

**Orthologous protein families**

The proteomes of eight mammals (Asiatic lion, Amur tiger, cat, dog, mouse, panda, opossum and human) were compared to examine the rate of protein evolution and conservation among protein orthologs using different databases meant for orthology prediction, OrthoMCL[54], InterPro[80] and Pfam[55].

**OrthoMCL:**

Among several databases and orthology prediction tools, the most sensitive and accurate is the OrthoMCL (utilizing the Markov Cluster Algorithm) program. OrthoMCL software is used to cluster proteins based on sequence similarity, using an all-against-all BLAST searchoff each species proteome, followed by normalization of inter-species differences, and Markov clustering. Based on sequence similarity and graph algorithm, OrthoMCL helps in distinguishing species proteomes into three separate categories; Orthologs, Co-orthologs and Paralogs, where the first two are speciation events (orthologsand co-orthologs) and the last one is the result of a duplication event (paralogs) within a genome(**Supplementary Table S21**).

The four felid proteomes of Asiatic lion, Amur tiger, domestic cat and Cheetah were compared to find Felidae-specific protein families. On combination of the four proteomes and removal of duplicates, a total of 126 protein families were found to be common to felids. Gene Ontology annotation could be done for 34 protein families (**Supplementary Table S24**). The PFam ID, InterPro ID and Ensembl ID were also determined for them. Comparisons with other genomes 17 genes found to be exclusive for lion and out of these 14 genes are annotated. The GO annotation, PFam ID, InterPro ID and Ensembl ID could be found for 9 protein families (**Supplementary Table S27**).

**Expansion-contraction of protein families and gain-loss of proteins:**

One method of assessing the phylogenetic relationships and deciphering the underlying evolutionary basis of certain significant metabolic phenotypes characterizing them is by evaluating the expansion-contraction of protein families or gain-loss of proteins among taxa.

Both these processes were examined by using the Wagner Parsimony algorithm of COUNT software computed by the AsymmetricWagner application[62] in this study.

COUNT is a software package for the analysis of numerical profiles on a phylogeny. COUNT performs ancestral reconstruction, and infers family- and lineage-specific characteristics along the evolutionary tree. It implements popular methods employed in gene content analysis such as Dollo[81] and Wagner parsimony[82], Propensity for Gene Loss[83], as well as probabilistic methods involving a phylogenetic birth-and-death model[84]. The asymmetric version of Wagner parsimony does differential penalization of gains and losses. Two input files were provided; the tree file (.txt format) and the Multiple Sequence Alignment (.tsv format) file. The output files were obtained for Dollo-Parsimony (DP), Wagner-Parsimony (WP) and PGL methods.

COUNT is for evolutionary analyses of phylogenetic profiles and other numerical characters and has been utilized here for estimating protein loss/gain and protein families' expansion/contraction. For ancestral reconstruction, asymmetric Wagner parsimony was applied and the gain penalty stringently set to 2.5. The lion protein families were further assigned to their InterPro IDs, GO and KEGG descriptions (**Supplementary Table S26**). PANTHER pathway analysis was also conducted (**Supplementary Table S27**).

**Positively selected genes:**

Detection of the strength and mode of natural selection on protein-coding genes is done by computing the $\omega$ statistic which is the ratio of rate of non-synonymous (dN) to synonymous (dS) substitutions[85]. $\omega > 1$ when positive selection occurs along multiple codons of a gene and throughout the entire phylogeny. But in most cases, positive selection is observed within a limited region of the protein and/or in some but not all the species in a phylogeny. Specific models have been developed to detect specific scenarios.

The statistic $\omega$ was estimated using the CODEML program of the Phylogenetic Analysis by Maximum Likelihood (PAML) package[56].dN and dS were calculated from orthologous protein-coding gene sequences obtained by alignment of annotated genomes of Asiatic lion, Amur tiger, cat, dog, mouse, panda, opossum and human. The orthologous gene sequences acted as input files for CODEML. The ortholog groups having more than one ortholog per species were removed retaining only 1:1 orthologs to avoid presence of paralogs that are a result of gene duplications. PRANK was used for multiple alignment of gene orthologs[57,58]. The branch-site model of PAML was used in which $\omega$ can vary at specific sites of a gene sequence and branches of the phylogeny[59]. A total of 240 lion protein-coding genes were determined as positively selected. The over-representation test was performed using the PANTHER database to identify gene families which are statistically over or under-represented in the list of positively selected genesbut none of the protein families were found to be significantly over-represented. A pathway analysis was also carried out using PANTHER for this list of genes **(Supplementary Table S28)**.

**Genomic diversity:**

The number of heterozygous SNVs per base pair of a genome has been used to comment on its genomic diversity in earlier papers on whole genomes[13,14]. The lion raw reads were mapped to the cat genome to determine the number of heterozygous SNVsin the lion genome

(**Supplementary Table S14**). The rate of heterozygous SNVs for 9 felid genomes was compiled from Kim *et al*.[13] and for lynx (Eurasian and Iberian) from Abascal *et al*.[19] to visualize the position of Asiatic lion among felid genomic diversity (**Supplementary Table S29**).

**Male-specific genes in lion genome:**

RNA-Seq dataof five Asiatic lion individuals (3 males, 2 females) were mapped to the lion genome using STAR aligner[86]. RNA-Seq by Expectation Maximization (RSEM) was used to quantitate and perform differential analysis with FDR < 0.5[87].Gene expression values (expected counts) were calculated using RSEM, where expected counts are the number of paired reads for each gene. Genes with two-fold change were considered as differentially enriched in male samples. Genes that are one-fold more enriched in males were listed as male-specific genes **(Supplementary Table S30).** Male-specific genes were selected based on a min value of 2 in PostFC which is the posterior fold change (condition 1 over condition 2) for a gene/transcript. It is defined as the ratio between posterior mean expression estimates of the gene/transcript for each condition.

**GENOME EVOLUTION:**

**Segmental duplications:**

The Whole Genome Assembly Comparison (WGAC) method was done to determine segmental duplications in the lion genome which is a BLAST based approach that performs an all-by-all comparison of assembled genomic sequence[59]. The method, however, makes two assumptions: (1) the genome has been accurately assembled and (2) no under- or over-representation within duplicated regions has occurred. A total of 6,174 recent segmental duplicated content of greater than 1kb length and more than 90% identity were found which accounts for 9.95Mb (0.4%) of the Asiatic lion genome. The low segmental duplication rate has been pointed out as an error in next generation sequencing assemblies by Cho *et al*[14] in the Amur tiger assembly.

**Chromosomal rearrangement:**

- The domestic cat (*Felis_catus* 8.0) and lion chromosomes (this study) were assembled using the program *nucmer* available from MUMmer[64] which is a package used for rapid alignment of entire genomes.
- When one region/block of a chromosome happened to be mapped to distant cat chromosomal locations, they were noted as inter- or intra-chromosomal rearrangement events of the lion genome relative to the cat genome.
- First, similar regions in intra-chromosomal comparisons were identified (Supplementary Figure L11). Here regions from same locations were discarded first. Then smaller regions which are less than 5Kb were discarded. Finally, the regions those are mapped to distant locations of the cat genome were listed.
- For inter-chromosomal comparisons, we considered all regions which are greater than 5Kb in length for rearrangement. Chromosomal rearrangement summary is shown in Supplementary Table S 35.
