## Supplemental Information for "De novo assembly and annotation of Asiatic lion (*Panthera leo persica*) genome"

- **Supplementary Figures**

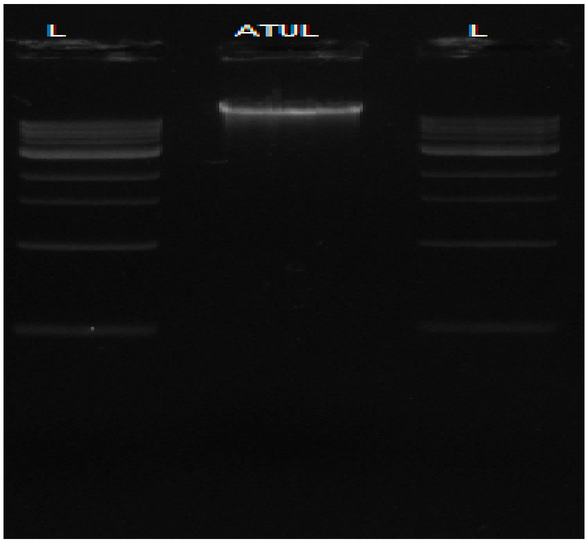


**Supplementary Figure L1: Agarose gel image of the genomic DNA isolated from AtuI**

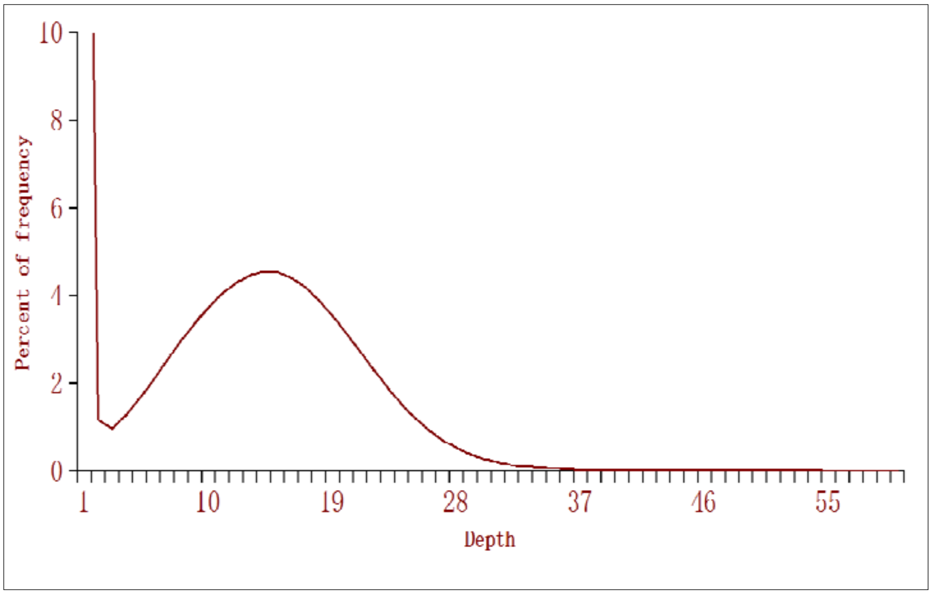


**Supplementary Figure L2: De Bruijn graph for genome size estimation using 23-mers**

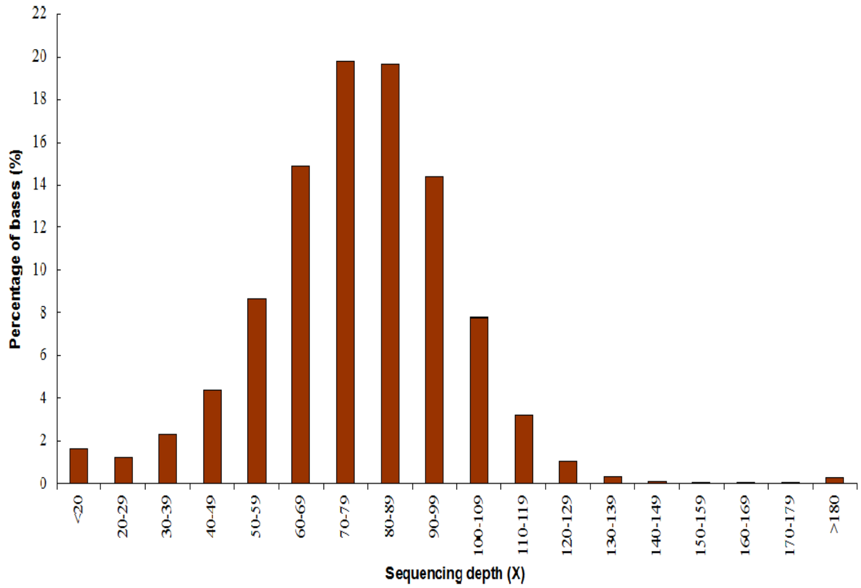


**Supplementary Figure L3: (A) Sequencing depth distribution**

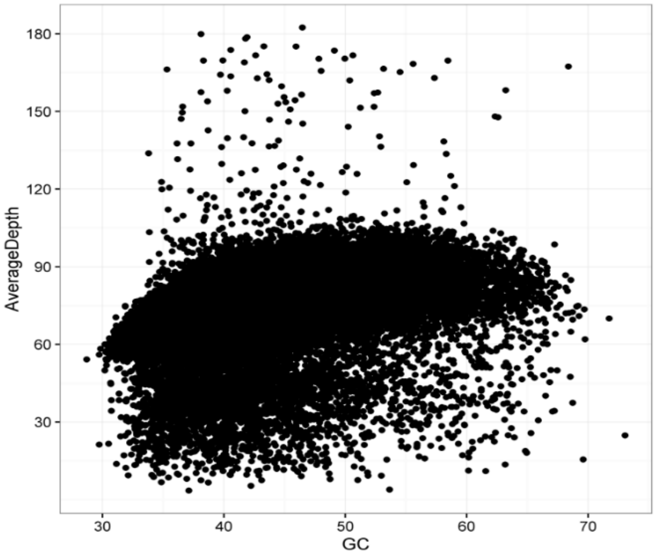


**Supplementary Figure L3 (B) GC contents and sequencing depths**

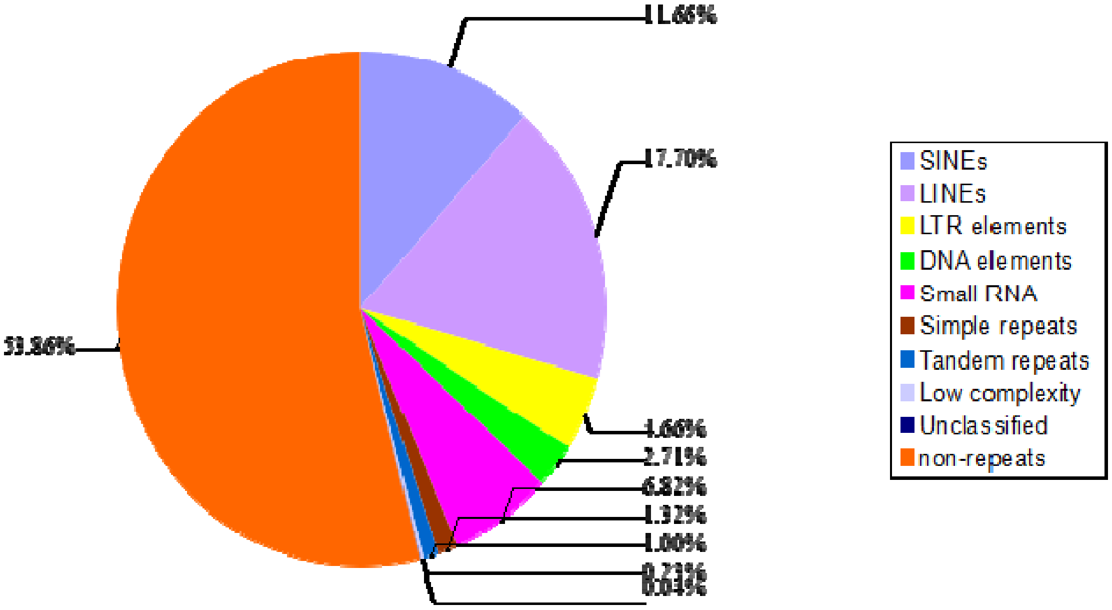


A

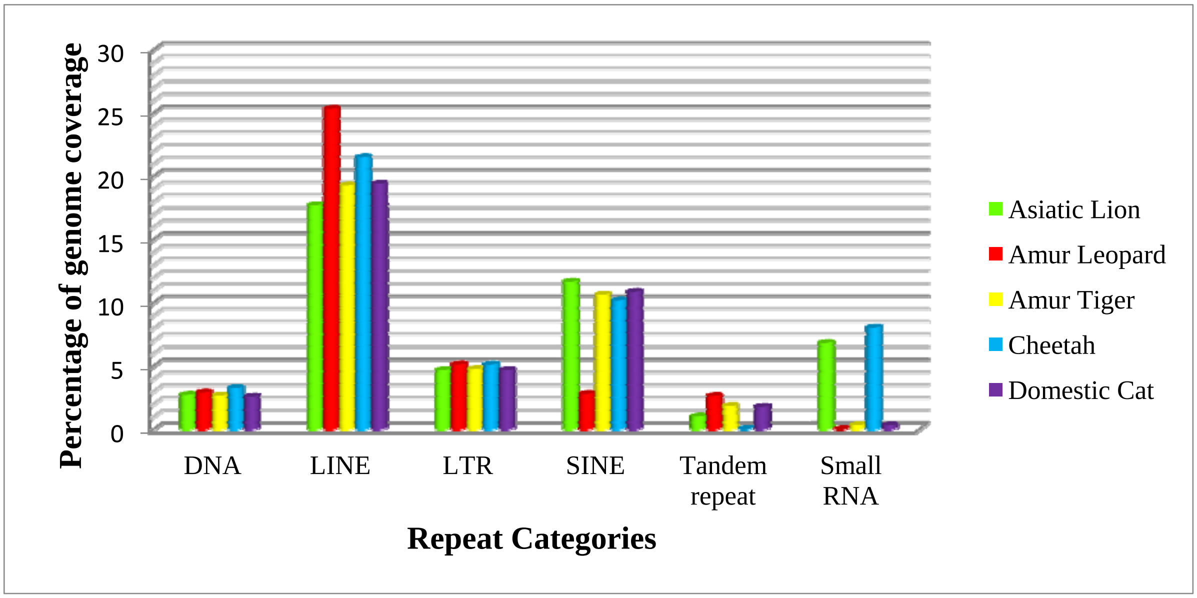


B

**Supplementary Figure L4: (A) Distribution of repeats in the Asiatic lion genome and (B) Comparison of repeats' distribution among Felidae genomes**

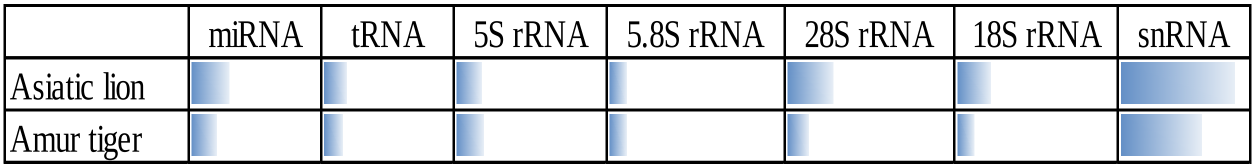


**Figure L5: Comparison of non-coding RNA with Amur tiger**

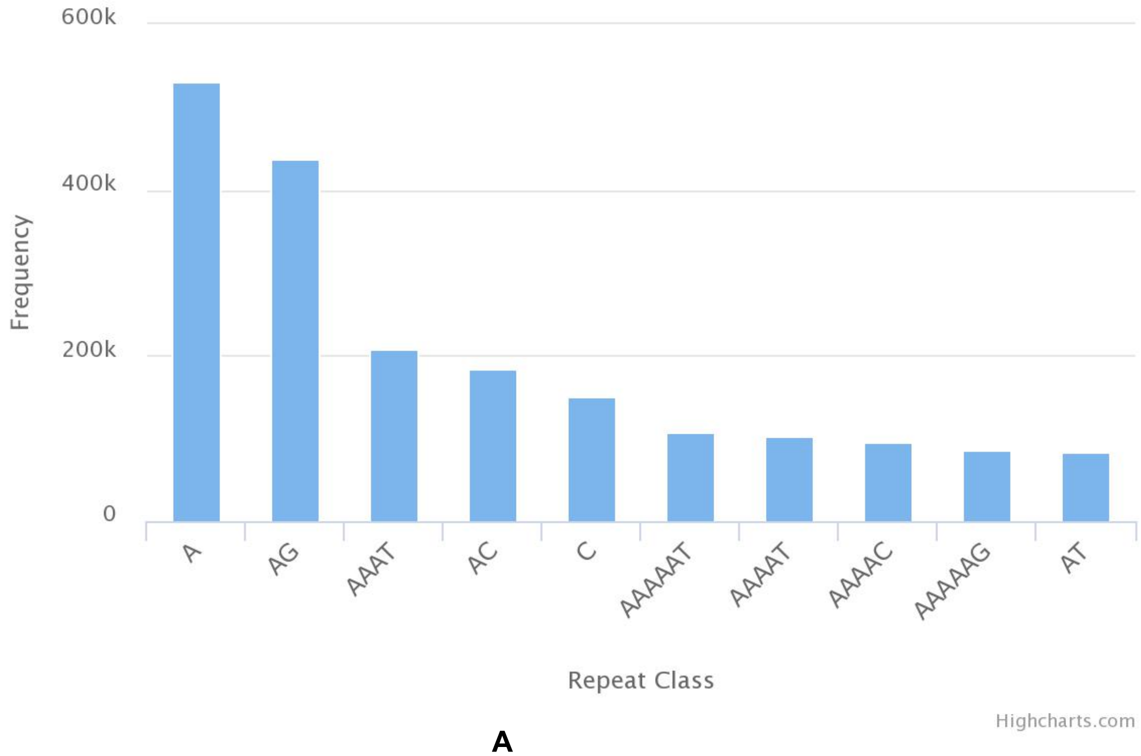


**A**

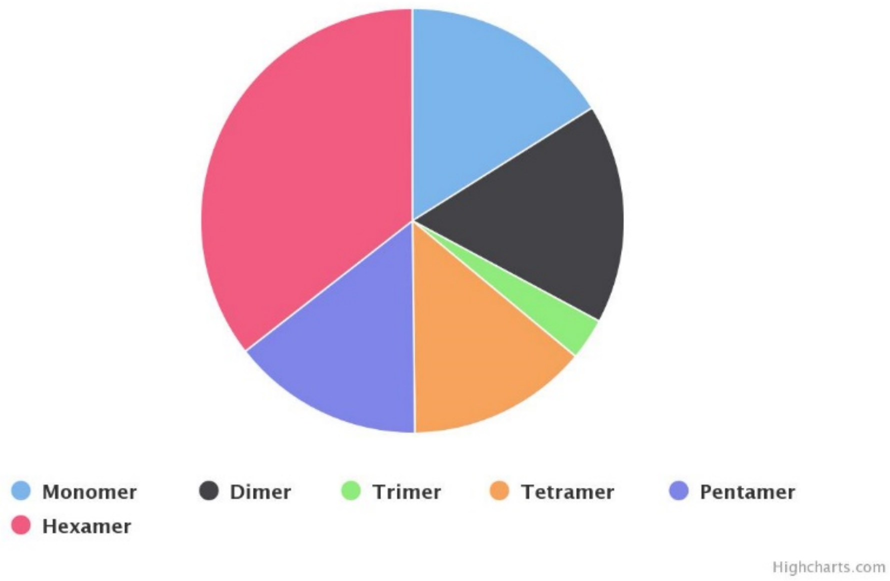


**Supplementary Figure L6: (A) Distribution of SSR repeats frequency in the Asiatic lion genome and (B) Distribution of SSRs**

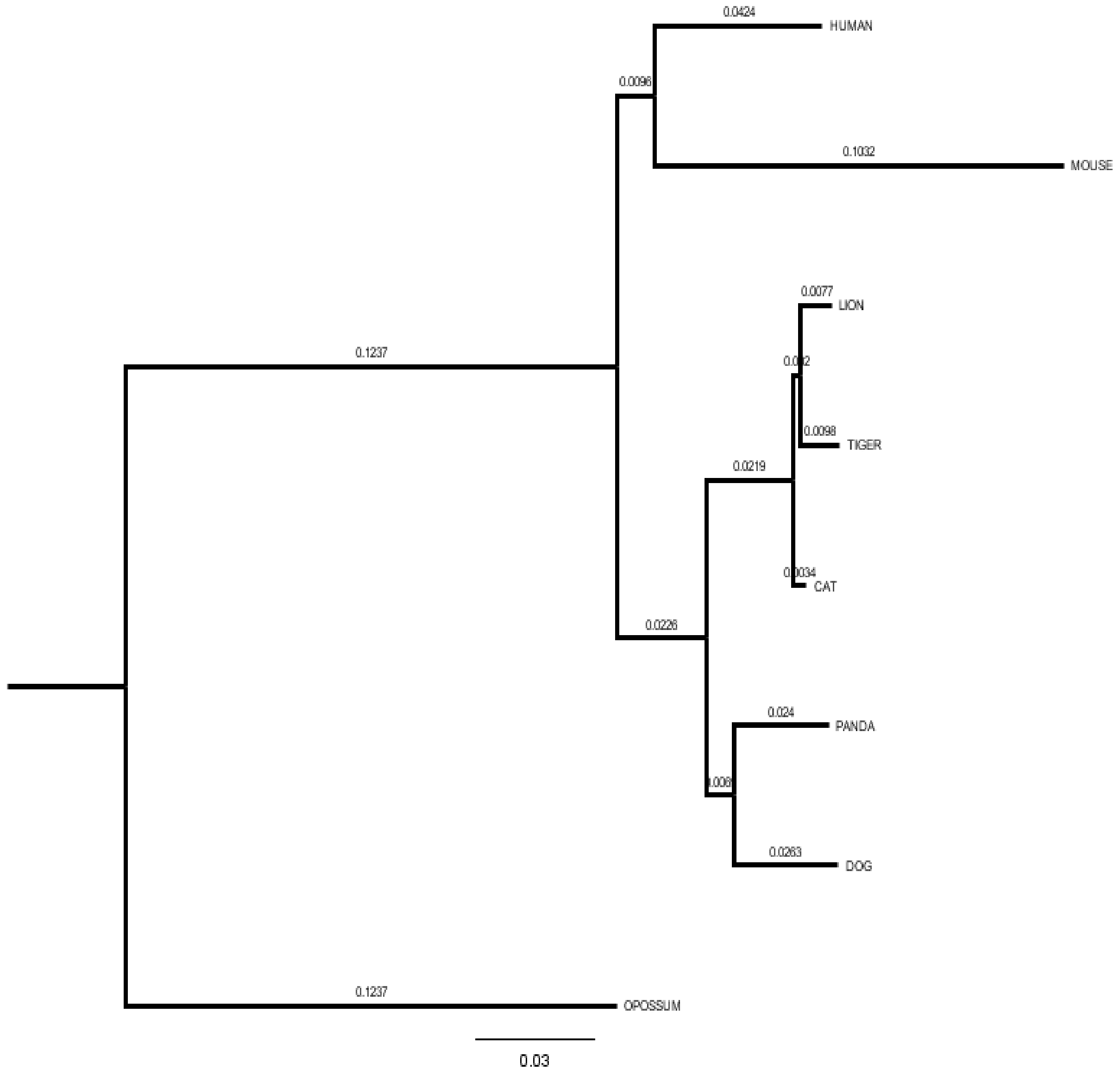


**Supplementary Figure L7: Maximum likelihood (ML) tree depicting the position of Asiatic lion in the mammalian phylogeny**

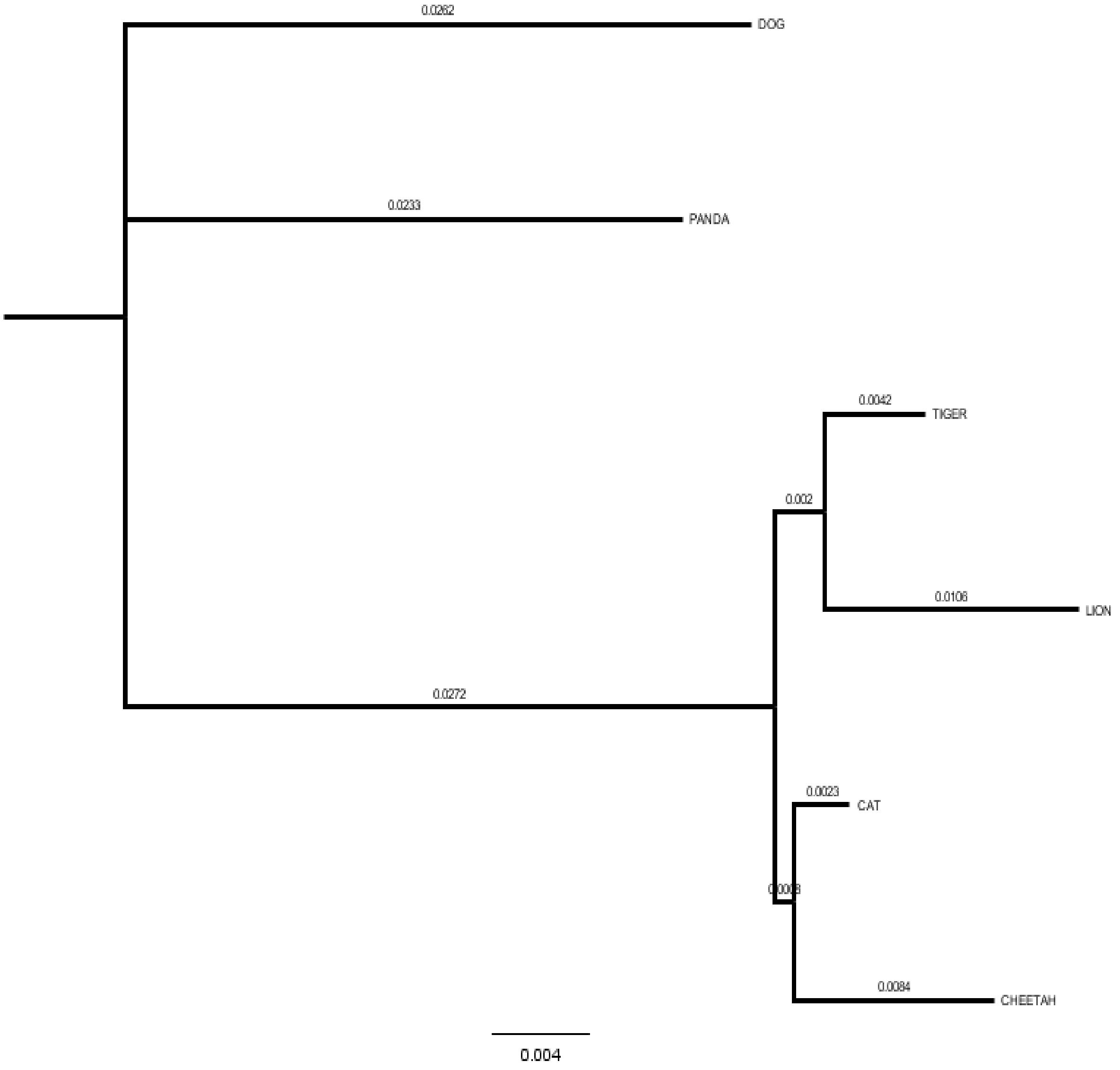


**Supplementary Figure L8: Maximum likelihood (ML) tree depicting the position of Asiatic lion in Carnivora phylogeny.**

**Supplementary Figure L9: Functional classification of positively selected genes in Asiatic lion into Biological Processes, Cellular Components and Molecular Functions**

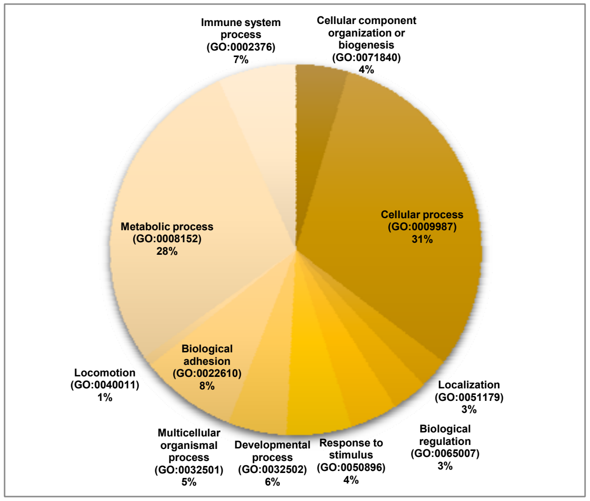


**Biological Processes**

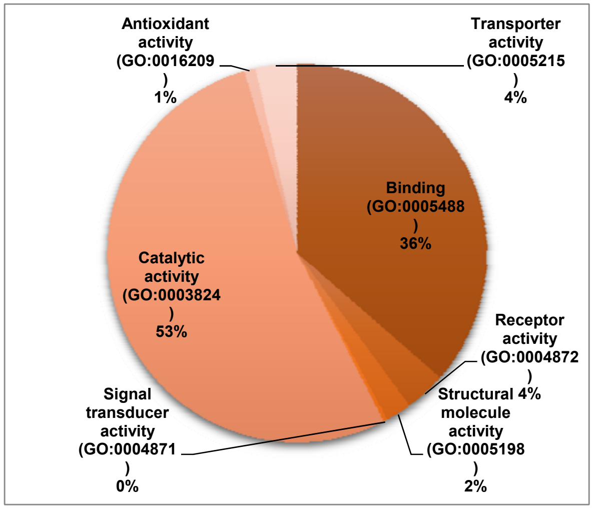


**Molecular Function**

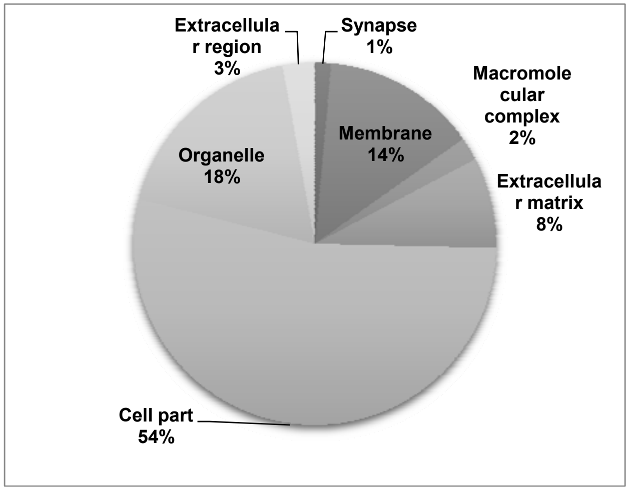


**Cellular components**

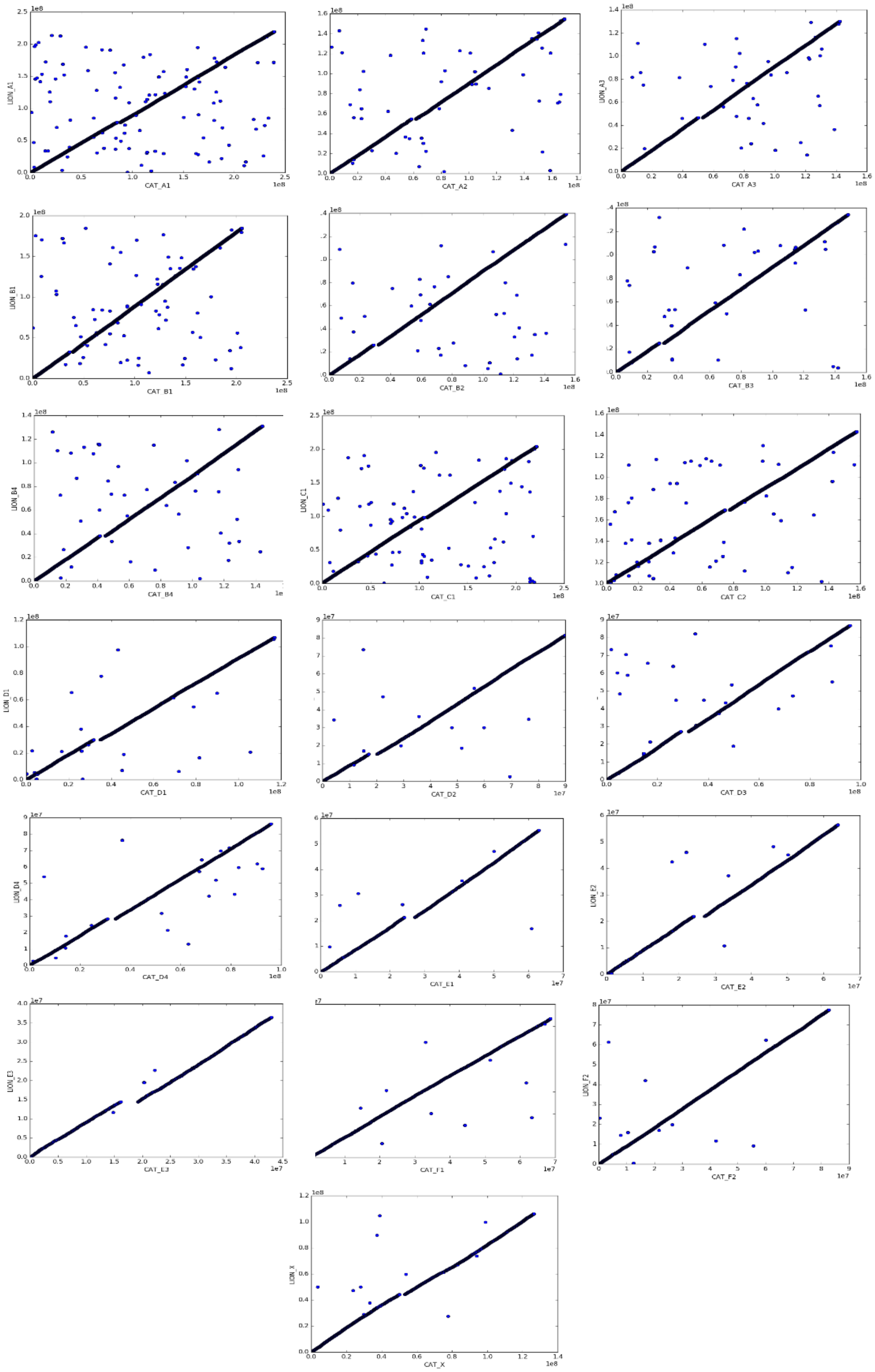


**Supplementary Figure L10: Dot plots or MUMs showing intra-chromosomal rearrangements with a block length cut off 5kb between Asiatic lion and domestic cat chromosomes. The x-axis represents the domestic cat genome (reference sequence) and the y-axis stands for the lion genome (query sequence). The line of blue dots shows exact conservation between two chromosomes. A disruption in this continuity is indicative of a rearrangement.**

### Supplementary Tables

#### Supplementary Table S1: Details of the blood sample used for sequencing

| Sample | Name | Sex | Place of Collection |
|---|---|---|---|
| Blood | **Atul** (National Stud Book No. 455) | Male | Nehru Zoological Park, Hyderabad |
| Blood | **Ajay** (National Stud Book No. 477) | Male | Nehru Zoological Park, Hyderabad |
| Blood | **Rita** (National Stud Book No. 454) | Female | Nehru Zoological Park, Hyderabad |
| Blood | **Soniya** (National Stud Book No. 572) | Female | Nehru Zoological Park, Hyderabad |
| Blood | **Vishwas** (National Stud Book No. 638) | Male | Nehru Zoological Park, Hyderabad |

**Supplementary Table S2: Quantification results of the extracted genomic DNA from blood sample of Atul**

| Sample Name | Qubit Reading conc. (ng/µl) | Nanodrop conc. (ng/µl) | 260/280 ratio |
|---|---|---|---|
| Atul | 161 | 318.9 | 1.8 |

**Supplementary Table S3: Description of the six pair-end and mate-pair libraries used for Asiatic lion genome assembly**

| Paired-end Libraries | Insert Size | Total Data (GB) | Average Read Length (bp) | Sequence Coverage(X) |
|---|---|---|---|---|
| Illumina Reads | 150bp | 55 | 100 | 22 |
| | 500bp | 42 | 100 | 17 |
| | 800bp | 24 | 100 | 10 |
| | 4-6Kb | 29 | 250 | 10 |
| | 6-10Kb | 25 | 250 | 8 |
| | 1-20Kb | 56 | 250 | 23 |
| **Total** | | **231** | | **90** |

**Supplementary Table S4: Information on the filtered sequence obtained**

| Paired-end libraries | Insert Size | Total Data(GB) | Average Read Length (bp) | Sequence Coverage(X) |
|---|---|---|---|---|
| Illumina Reads | 150bp | 48 | 100 | 20 |
| | 500bp | 35 | 100 | 14 |
| | 800bp | 18 | 100 | 7 |
| | 4-6Kb | 12 | 250 | 5 |
| | 6-10Kb | 12 | 250 | 5 |
| | 1-20Kb | 27 | 250 | 11 |
| **Total** | | **152** | | **62** |

**Supplementary Table S5: Genome sizes obtained by using K-mer values 17, 21 and 23**

| K | Genome size (in bp) |
|---|---|
| 17 | 1,240,073,836 |
| 21 | 1,951,822,091 |
| 23 | 2,033,113,726 |

**Supplementary Table S6: Statistics for checking assembly quality**

| | Size (bp) | Contig Number | Size (bp) | Scaffold Number |
|---|---|---|---|---|
| **N90** | 2,900 | 223,929 | 14,580 | 63,620 |
| **N80** | 4,700 | 163,455 | 19,315 | 48,654 |
| **N70** | 6,620 | 118,529 | 24,685 | 37,308 |
| **N60** | 8,545 | 87,453 | 30,300 | 28,308 |
| **N50** | 10,669 | 62,599 | 36,647 | 20,864 |
| **Longest** | 143,273 | - | 383,982 | - |
| **Total Size** | 2,365,521,145 | - | 2,461,536,221 | - |
| **Total Number(>1000bp)** | 2,316,348,500 | 323,968 | 2,452,616,604 | 94,372 |
| **Total Number(>5Kb)** | 1,866,770,627 | 155,481 | 2,427,573,497 | 83,167 |

**Supplementary Table S7: Asiatic lion blood transcriptomes statistics**

| Individual's Name | Number of total reads | Total nucleotides (nt) | > Q20 (%) | N base (%) | GC contents (%) |
|---|---|---|---|---|---|
| Ajay | 87,178,978 | 17,435,795,600 | 91.04 | 0.0768 | 48.42 |
| Atul | 51,348,948 | 10,269,789,600 | 88.74 | 0.0514 | 51.38 |
| Rita | 48,860,522 | 9,772,104,400 | 90.36 | 0.0717 | 45.56 |
| Sonia | 60,782,750 | 12,156,550,000 | 86.69 | 0.0732 | 51.72 |
| Viswas | 98,594,244 | 19,718,848,800 | 89.96 | 0.0870 | 49.12 |

**Supplementary Table S8: Gene coverage evaluation from assembled Asiatic lion transcripts**

| Data set | Number | Total Length (bp) | Covered by Assembly | With >90% sequence in one scaffold | | With >50% sequence in one scaffold | |
|---|---|---|---|---|---|---|---|
| | | | | Number | Percent (%) | Number | Percent (%) |
| All | 1,519,892 | 787,784,760 | 81.78 | 901,883 | 59.33 | 1,304,378 | 85.82 |
| >200 | 1,519,892 | 787,784,760 | 81.78 | 901,883 | 59.33 | 1,304,378 | 85.82 |
| >500 | 373,840 | 447,253,979 | 81.33 | 201,352 | 53.86 | 323,994 | 86.66 |
| >1000 | 142,537 | 290,521,816 | 81.86 | 73,674 | 51.68 | 127,114 | 89.17 |

**Supplementary Table S9: Gene coverage evaluation from assembled Asiatic lion transcripts**

| Total EST | Mapped EST | Mapped Percent (%) | EST s Covered by Assembly (%) | >90% mapped in one scaffold | | >50% mapped in one scaffold | | >20% mapped in one scaffold | |
|---|---|---|---|---|---|---|---|---|---|
| | | | | Number | Percent | Number | Percent | Number | Percent |
| 919 | 888 | 96.63 | 86.86 | 630 | 68.55 | 820 | 89.23 | 865 | 94.12 |

**Supplementary Table S10: Evolution of the completeness of the Asiatic lion genome assembly using (CEGMA) approach.**

| **Parameter** | **Number** | **Percent (%)** |
|---|---|---|
| Total KOGs | 458 | |
| ONE KOG align one gene | 391 | 86.89 |
| ONE KOG align no gene | 67 | 14.62 |

**Supplementary Table S11: Comparison of Asiatic lion and domestic cat genomes (Felis_catus-8.0) showing number of chromosomes, base composition and GC content.**

| | Lion | | Domestic cat | |
|---|---|---|---|---|
| **Number of chromosomes** | 19 | | 19 | |
| **Total bases** | 2,119,661,238 | | 2,428,540,393 | |
| **Base** | **Number** | **Percentage** | **Number** | **Percentage** |
| **A** | 589,629,285 | 27.82 | 681,725,917 | 28.07 |
| **T** | 590,066,774 | 27.84 | 682,210,425 | 28.09 |
| **G** | 420,116,318 | 19.82 | 487,148,479 | 20.06 |
| **C** | 419,941,851 | 19.81 | 486,985,203 | 20.05 |
| **N** | 99,907,010 | 4.71 | 90,470,369 | 3.73 |
| **GC Content (%)** | 39.63% | | 41.66% | |

**Supplementary Table S12: Statistical figures obtained by mapping Asiatic lion raw reads to the domestic cat genome (*Felis_catus*-8.0)**

| | Total bases (except N) | No depth filter | ≥ 5 Depth | Coverage (no depth filter) | Coverage (up to 5 depth) |
|---|---|---|---|---|---|
| ChrA1 | 239,302,903 | 230,898,223 | 229,240,912 | 0.9649 | 0.9580 |
| ChrA2 | 169,043,629 | 161,904,742 | 160,691,623 | 0.9578 | 0.9506 |
| ChrA3 | 142,459,683 | 136,304,895 | 135,254,076 | 0.9568 | 0.9494 |
| ChrB1 | 205,241,052 | 196,811,928 | 195,144,805 | 0.9589 | 0.9508 |
| ChrB2 | 154,261,789 | 147,133,649 | 145,873,840 | 0.9538 | 0.9456 |
| ChrB3 | 148,491,654 | 141,273,585 | 140,046,675 | 0.9514 | 0.9431 |
| ChrB4 | 144,259,557 | 137,335,845 | 136,184,096 | 0.9520 | 0.9440 |
| ChrC1 | 221,441,202 | 213,110,793 | 211,477,448 | 0.9624 | 0.9550 |
| ChrC2 | 157,659,299 | 151,017,716 | 149,831,156 | 0.9579 | 0.9503 |
| ChrD1 | 116,869,131 | 110,726,288 | 109,793,732 | 0.9474 | 0.9395 |
| ChrD2 | 89,822,065 | 84,566,967 | 83,896,584 | 0.9415 | 0.9340 |
| ChrD3 | 95,741,729 | 90,320,473 | 89,604,304 | 0.9434 | 0.9359 |
| ChrD4 | 96,020,406 | 90,589,512 | 89,856,654 | 0.9434 | 0.9358 |
| ChrE1 | 63,002,102 | 58,128,433 | 57,624,952 | 0.9226 | 0.9147 |
| ChrE2 | 64,039,838 | 59,012,562 | 58,491,914 | 0.9215 | 0.9134 |
| ChrE3 | 43,024,555 | 38,842,709 | 38,516,971 | 0.9028 | 0.8952 |
| ChrF1 | 68,669,167 | 66,702,383 | 66,165,940 | 0.9714 | 0.9635 |
| ChrF2 | 82,763,536 | 80,822,390 | 80,204,363 | 0.9765 | 0.9691 |
| ChrX | 126,427,096 | 115,457,488 | 113,206,614 | 0.9132 | 0.8954 |
| Avg. | - | - | - | 0.9473 | 0.9391 |

**Supplementary Table S13: Statistics regarding mapping of Asiatic lion raw reads to the Asiatic lion scaffolds**

| Total reads | Mapped reads | % of mapped reads | Properly mapped reads | Unmapped | Coverage |
|---|---|---|---|---|---|
| 1,588,787,434 | 1,535,593,778 | 96.65 | 1,386,737,899 | 53,196,656 | 55.5 |

**Supplementary Table S14: Variation Statistics regarding mapping of lion reads to the Asiatic lion scaffolds**

| | Homozygous | Heterozygous | Total |
|---|---|---|---|
| **SNVs** | 221,397 | 22,790 | 244,187 |
| **INDELs** | 39,860 | 15,727 | 55,587 |
| **Total** | 261,257 | 38,517 | 299,774 |

**Supplementary Table S15: Variation Statistics regarding mapping of Asiatic lion reads to the domestic cat genome (*Felis_catus-8.0*)**

| | **Homozygous** | **Heterozygous** | **Total** |
|---|---|---|---|
| **SNV s** | 41,532,416 | 745,184 | 42,277,600 |
| **INDELs** | 6,980,946 | 533,005 | 7,513,951 |
| **Total** | 48,513,362 | 1,278,189 | 49,791,551 |

**Supplementary Table S16: Statistics of Asiatic lion draft chromosomes**

| | Total bases | Number of A, C, G, T | Number of N | Number of scaffolds | Avg. length of scaffolds | Max. length of scaffold | Min. length of scaffold |
|---|---|---|---|---|---|---|---|
| **ChrA1** | 212,374,592 | 204,199,597 | 8,174,995 | 11,557 | 21,207 | 205,056 | 88 |
| **ChrA2** | 149,495,056 | 142,948,854 | 6,546,202 | 8,353 | 20,595 | 262,985 | 78 |
| **ChrA3** | 125,695,176 | 120,729,876 | 4,965,300 | 6,677 | 21,790 | 267,142 | 85 |
| **ChrB1** | 178,427,797 | 170,735,598 | 7,692,199 | 10,218 | 20,111 | 201,034 | 54 |
| **ChrB2** | 134,423,980 | 128,830,742 | 5,593,238 | 7,594 | 20,499 | 195,532 | 96 |
| **ChrB3** | 129,299,323 | 123,734,213 | 5,565,110 | 7,306 | 20,576 | 214,589 | 96 |
| **ChrB4** | 126,213,181 | 120,752,019 | 5,461,162 | 7,155 | 20,432 | 246,013 | 65 |
| **ChrC1** | 197,424,461 | 189,172,046 | 8,252,415 | 10,734 | 21,100 | 229,378 | 71 |
| **ChrC2** | 138,521,516 | 132,845,619 | 5,675,897 | 7,633 | 20,855 | 383,982 | 93 |
| **ChrD1** | 103,188,731 | 98,421,593 | 4,767,138 | 6,080 | 19,370 | 216,203 | 96 |
| **ChrD2** | 78,837,955 | 75,633,457 | 3,204,498 | 4,186 | 21,755 | 214,418 | 103 |
| **ChrD3** | 83,982,566 | 80,474,991 | 3,507,575 | 4,549 | 21,445 | 233,055 | 90 |
| **ChrD4** | 83,215,589 | 79,456,265 | 3,759,324 | 4,669 | 20,667 | 240,205 | 72 |
| **ChrE1** | 52,862,329 | 49,744,235 | 3,118,094 | 3,170 | 19,613 | 205,393 | 61 |
| **ChrE2** | 54,133,435 | 51,299,689 | 2,833,746 | 3,085 | 20,502 | 209,317 | 60 |
| **ChrE3** | 35,050,219 | 33,302,104 | 1,748,115 | 2,050 | 19,899 | 207,721 | 95 |
| **ChrF1** | 60,922,062 | 58,301,526 | 2,620,536 | 3,638 | 19,441 | 170,669 | 64 |
| **ChrF2** | 74,891,923 | 72,135,184 | 2,756,739 | 4,074 | 21,133 | 205,397 | 92 |
| **ChrX** | 100,701,347 | 87,036,620 | 13664727 | 9,463 | 11,917 | 146,549 | 111 |
| **Total** | 2,119,701,455 | 2,019,782,242 | 99,919,213 | 122,195 | - | - | - |

**Supplementary Table S17: Summary of non-coding RNA in Asiatic lion genome:**

| Type | | Copy number | Average length (bp) | Total length (bp) | % of genome |
|---|---|---|---|---|---|
| **miRNA** | | 1010 | 74 | 75,377 | 0.00305 |
| **tRNA** | | 413 | 74 | 30,562 | 0.00123 |
| **rRNA** | 5S | 234 | 112 | 26,200 | 0.00106 |
| | 5.8S | 4 | 153 | 613 | 0.00002 |
| | 18S rRNA | 75 | 895 | 67,141 | 0.00271 |
| | 28S rRNA | 37 | 1,084 | 40,110 | 0.00162 |
| **snRNA** | snRNA total, spliceosomal RNA, CD-box | 2,284 | 134 | 3,07,887 | 0.01246 |

**Supplementary Table S18: Statistics of predicted protein-coding genes in Asiatic lion as compared to other genomes using various models**

| | Gene set | Number | Average transcript length (bp) | Average CDS length (bp) | Average no. of exons per gene | Average exons length (bp) | Average intron length |
|---|---|---|---|---|---|---|---|
| *De novo* | **AUGUSTUS** | 19,086 | 23,860 | 1,097 | 7 | 161 | 3,287 |
| | **Cat** | 19,945 | 10,786 | 1,240 | 7 | 190 | 1,724 |
| | **Tiger** | 18,943 | 10,871 | 1,196 | 6 | 186 | 1,780 |
| **Homology** | **Dog** | 20,362 | 11,337 | 1,264 | 7 | 187 | 1,749 |
| | **Human** | 28,873 | 11,720 | 1,210 | 7 | 175 | 1,777 |
| | **Mouse** | 23,613 | 11,356 | 1,224 | 7 | 180 | 1,742 |
| **EST** | | 719 | 1,391 | 512 | 3 | 177 | 465 |
| **EVM** | | 20,543 | 13,818 | 1,083 | 7 | 162 | 2,391 |

**Supplementary Table S19: Statistics of predicted protein-coding genes in Asiatic lion as compared to other genomes: Evidence from EVM gene models**

| | | ≥ 20% overlap | | ≥ 50% overlap | | ≥ 80% overlap | |
|---|---|---|---|---|---|---|---|
| | | **Number** | **Percent** | **Number** | **Percent** | **Number** | **Percent** |
| **Augustus** | | 17,980 | 87.52 | 16,798 | 81.77 | 14,173 | 68.99 |
| | **Cat** | 8,186 | 39.85 | 3,402 | 16.56 | 1,503 | 7.32 |
| | **Tiger** | 7,903 | 38.47 | 3,222 | 15.68 | 1,408 | 6.85 |
| **Exonerate** | **Dog** | 7,503 | 36.52 | 2,995 | 14.58 | 1,300 | 6.33 |
| | **Human** | 7,656 | 37.27 | 2,918 | 14.20 | 1,190 | 5.79 |
| | **Mouse** | 6,962 | 33.89 | 2,694 | 13.11 | 1,136 | 5.53 |
| **EST** | | 167 | 0.81 | 80 | 0.39 | 37 | 0.18 |

**Supplementary Table 20: Number of genes in Asiatic lion annotated for homology or functional classification by CANoPI**

| | | Number | Percent |
|---|---|---|---|
| **Total** | | 20,543 | - |
| **Annotated** | InterPro | 19,316 | 94.03 |
| | GO | 17,277 | 84.10 |
| | KEGG | 4,543 | 22.11 |
| | SwissProt | 178 | 0.87 |
| | TrEMBL | 19,825 | 96.50 |
| **Unannotated** | | 540 | 2.63 |

**Supplementary Table S21: Statistical analysis of orthologous protein families in Asiatic lion using PFam**

| Species | Single copy orthologs | Co-orthologs | Unique paralogs | Other orthologs | Un-clustered | Total |
|---|---|---|---|---|---|---|
| **Asiatic lion** | **6,295** | **4** | **21** | **8,024** | **6,199** | **20,543** |
| Cat | 9,447 | 59 | 100 | 22,654 | 960 | 33,220 |
| Amur tiger | 8,806 | 41 | 110 | 20,516 | 0 | 29,473 |
| Human | 9,591 | 1,621 | 4,768 | 82,150 | 0 | 98,130 |
| Mouse | 9,755 | 1,357 | 4,001 | 61,028 | 0 | 76,141 |
| Dog | 5,846 | 163 | 311 | 40,767 | 0 | 47,087 |
| Panda | 13,810 | 472 | 694 | 17,531 | 0 | 32,507 |
| Opossum | 10,672 | 1,378 | 4,218 | 32,844 | 0 | 49,112 |

**Supplementary Table S22: Functional classification of Felidae-specific protein families**

| Database* | Description | GO term | Number of genes |
|---|---|---|---|
| BP | Cellular component organization or biogenesis | GO:0071840 | 8 |
| | Cellular process | GO:0009987 | 24 |
| | Localization | GO:0051179 | 6 |
| | Reproduction | GO:0000003 | 1 |
| | Biological regulation | GO:0065007 | 6 |
| | Response to stimulus | GO:0050896 | 4 |
| | Developmental process | GO:0032502 | 2 |
| | Multicellular organismal process | GO:0032501 | 1 |
| | Biological adhesion | GO:0022610 | 1 |
| | Metabolic process | GO:0008152 | 24 |
| | Immune system process | GO:0002376 | 4 |
| MF | Binding | GO:0005488 | 18 |
| | Receptor activity | GO:0004872 | 1 |
| | Structural molecule activity | GO:0005198 | 3 |
| | Catalytic activity | GO:0003824 | 14 |
| | Transporter activity | GO:0005215 | 4 |
| CC | Membrane | GO:0016020 | 1 |
| | Macromolecular complex | GO:0032991 | 8 |
| | Cell part | GO:0044464 | 17 |
| | Organelle | GO:0043226 | 11 |
| | Extracellular region | GO:0005576 | 3 |

***BP: Biological Process, MF: Molecular Function, CC: Cellular Components**

**Supplementary Table S23: PANTHER pathway analysis of Felidae-specific protein families**

| **PANTHER Pathway** | **Total Components** | **Felidae Genes** |
|---|---|---|
| BMP/activin signaling pathway-drosophila | 37 | 1 |
| JAK/STAT signaling pathway | 19 | 2 |
| B cell activation | 36 | 1 |
| Interleukin signaling pathway | 21 | 2 |
| Interferon-gamma signaling pathway | 3 | 1 |
| Inflammation mediated by chemokine and cytokine signaling pathway | 28 | 1 |
| Asparagine and aspartate biosynthesis | 8 | 1 |
| T cell activation | 18 | 1 |
| TGF-beta signaling pathway | 36 | 1 |
| SCW signaling pathway | 45 | 1 |
| PDGF signaling pathway | 20 | 1 |
| EGF receptor signaling pathway | 19 | 1 |
| DPP signaling pathway | 58 | 1 |
| DPP-SCW signaling pathway | 10 | 1 |

**Supplementary Table S24: Gene Ontology ID, GO terms and InterPro IDs of Felidae-specific protein families**

| PFAM ID | InterPro ID | TYPE | GO Term |
|---|---|---|---|
| PF00174.17 | IPR000572 | Domain | Nitrate assimilation |
| PF00190.20 | IPR006045 | Domain | Nutrient reservoir activity |
| PF00363.16 | IPR001588 | Family | Transporter activity, transport, extracellular region |
| PF00665.24 | IPR001584 | Domain | DNA Integration |
| PF00714.15 | IPR002069 | Family | Interferon-gamma receptor binding, immune response, extracellular region |
| PF01958.16 | IPR002811 | Domain | Oxidoreductase activity, NADP catabolic process, pyridine nucleotide biosynthetic process, oxidation-reduction process |
| PF02093.14 | IPR003036 | Domain | Virion assembly |
| PF02545.12 | IPR003697 | Family | Cytoplasm |
| PF02765.15 | IPR011564 | Domain | DNA binding, telomere maintenance, nuclear chromosome |
| PF03404.14 | IPR005066 | Domain | Oxidoreductase activity, molybdenum ion binding, oxidation-reduction process |
| PF04030.12 | IPR007173 | Domain | D-arabinono-1,4-lactone oxidase activity, oxidation-reduction process, membrane |
| PF04420.12 | IPR028945 | Family | Tail-anchored membrane protein insertion into ER membrane |
| PF05669.10 | IPR008831 | Family | RNA polymerase II transcription cofactor activity, regulation of transcription DNA-templated, mediator complex |
| PF05808.9 | IPR008783 | Family | Integral component of membrane |
| PF06214.9 | IPR010407 | Domain | Receptor activity, lymphocyte activation, cell surface, integral component of membrane |
| PF07855.10 | IPR012445 | Family | Autophagy |
| PF07994.10 | IPR002587 | Family | Inositol-3-phosphate synthase activity, inositol biosynthetic process, phospholipid biosynthetic process |
| PF08451.9 | IPR013659 | Domain | Extracellular space |
| PF09236.8 | IPR015317 | Family | Hemoglobin binding, protein folding, erythrocyte differentiation, protein |

| | | | |
|---|---|---|---|
| | | | stabilization |
| PF09252.8 | IPR015332 | Family | Extracellular space |
| PF09412.8 | IPR018998 | Family | Hydrolase activity acting on ester bonds |
| PF10204.7 | IPR018469 | Family | Protein transport, endoplasmic reticulum membrane, integral component of membrane |
| PF10461.7 | IPR019502 | Family | Apoptotic process, cellular response to DNA damage stimulus |
| PF10471.7 | IPR018860 | Family | Regulation of mitotic metaphase/anaphase transition, anaphase-promoting complex-dependent catabolic process, anaphase-promoting complex |
| PF12567.6 | IPR016335 | Family | Protein tyrosine phosphatase activity, T cell receptor signaling pathway |
| PF12632.5 | IPR026859 | Domain | Myosin binding |
| PF13096.4 | IPR027801 | Family | CENP-A containing nucleosome assembly, chromosome, centromeric region |
| PF15077.4 | IPR027816 | Family | DNA binding |
| PF15307.4 | IPR029301 | Family | Acrosomal vesicle |
| PF15510.4 | IPR028847 | Family | DNA binding, mitotic nuclear division, kinetochore assembly |
| PF15549.4 | IPR029096 | Family | Methylated histone binding |
| PF15677.3 | IPR020162 | Family | Cerebellum development, neuron differentiation |
| PF15703.3 | IPR031428 | Family | Immune response-regulating signaling pathway, B cell activation |
| PF15718.3 | IPR031447 | Family | Centriole replication |

**Supplementary Table S25: Gene Ontology terms and InterPro IDs of Asiatic lion-specific protein families**

| InterPro | TYPE | GO term |
|---|---|---|
| IPR015872 | Domain | Transcription initiation from RNA polymerase II promoter, transcription factor TFIIA complex |
| IPR015871 | Domain | Transcription initiation from RNA polymerase II promoter, transcription factor TFIIA complex |
| IPR004850 | Domain | Laminin binding, G-protein coupled acetylcholine receptor signaling pathway, receptor clustering |
| IPR005106 | Domain | Oxidoreductase activity, NADP binding, oxidation-reduction process |
| IPR005314 | Family | Peptidase activity, proteolysis, nucleus |
| IPR007747 | Family | Nucleus |
| IPR009395 | Family | BLOC-1 complex |
| IPR013221 | Domain | ATP binding, biosynthetic process |
| IPR028067 | Family | Immune response |

**Supplementary Table S26: Protein-families that underwent expansion in Asiatic lions**

| Protein family | InterPro ID | GO ID Description | KEGG ID Description |
|---|---|---|---|
| AICARFT_IMPCHas | IPR002695 | | KO0230Purine metabolism; KO0670one carbon pool by folate |
| Aquarius_N | IPR032174 | 0005681 Spliceosomal complex; 0000398mRNA splicing, via spliceosome | KO12874Genetic information processing |
| CDO_I | IPR010300 | 0017172cysteine dioxygenase activity; 0005506iron ion binding, 0055114oxidation-reduction process;0046439 L-cysteine metabolic process | KO0270Cysteine and methionine metabolism; KO0430 taurine and hypotaurine metabolism |
| CLP_protease | IPR001907 | | KO04112cell cycle; KO04212longevity regulating pathway |
| Cys_rich_FGFR | IPR001893 | 0016020 | |
| DNA_pol_phi | IPR007015 | 0003887DNA-directed DNA polymerase activity; 0003677DNA binding;0006351DNA templated transcription | KO02331 |
| DRIM | IPR011430 | | KO14772ribosome biogenesis |
| DUF2046 | IPR019152 | | KO05200Pathways in cancer, KO05216thyroid cancer |
| eIF3g | IPR024675 | | KO03013RNA transport |
| eIF-3_zeta | IPR007783 | 0005852Eukaryotic translation initiation factor 3 complex; 0005737cytoplasm; 0003743translation initiation factor activity | KO03013RNA transport |
| Gaa1 | IPR007246 | 0042765GPI-anchor transamidase complex; 0016021integral component of membrane | KO00563Glycosylphosphatidylinositol (GPI)-anchor |

| | | | |
|---|---|---|---|
| | | | biosynthesis |
| Lyase_aromatic | IPR001106 | | KO00340Histidine metabolism |
| NUC202 | IPR012980 | | KO03009ribosome biogenesis |
| OSTMP1 | IPR019172 | | |
| PRP1_N | IPR010491 | 0005634 Nucleus, 0000398mRNA splicing, via spliceosome | KO03040Spliceosome |
| Ribosomal_L23eN | IPR005633 | | KO03010Ribosome |
| Ribosomal_S26e | IPR000892 | 0005840Ribosome; 0005622intracellular; 0003735 Structural constituents of ribosome; 0006412 Translation | KO03010Ribosome |
| SBP_bac_3 | IPR001638 | | KO02030 |
| SKIP_SNW | IPR004015 | 0005681spliceosomal complex; 0000398mRNA splicing, via spliceosome | KO03040Spliceosome; KO04330notch signalling pathway; KO05169epstein-barr virus infection; KO05203viral carcinogenesis |
| SRP68 | IPR026258 | 0005786 Signal Recognition Particle, endoplasmic reticulum targeting;0030942 endoplasmic reticulum signal peptide binding;0005047 signal recognition particle binding; 00083127S RNA binding; 0006614SRP dependent co-translational protein targeting to membrane | KO03060Protein export |
| TIM | IPR000652 | 0004807triose phosphate isomerase activity; 0008152metabolic process | KO00010glycolysis/gluconeogenesis; KO00051fructose and mannose metabolism; KO00562 inositol phosphate metabolism; KO00710carbon |

| | | | |
|---|---|---|---|
| | | | metabolism; KO01200amino acid biosynthesis |
| TMA7 | IPR015157 | | |
| U5_2-snRNA_bdg | IPR019581 | 0030623U5 snRNA binding | KO03040Spliceosome |
| WT1 | IPR000976 | 0005634 Nucleus; 0006355 Regulation of transcription, DNA templated | KO05202Transcriptional misregulation in cancer |

**Supplementary Table S27: PANTHER pathway analysis of protein families expanded in Asiatic lions**

| PANTHER Pathway | Pathway ID | Total Components | Asiatic Lion Genes | Fold over-represented | *p*-value |
|---|---|---|---|---|---|
| Ionotropic glutamate receptor pathway | PTHR18966:SF269 | 29 | 1 | 36.68 | 0 |
| De novo purine biosynthesis | PTHR11692:SF3 | 23 | 1 | 26.90 | 0 |
| Metabotropic glutamate receptor group III pathway | PTHR18966:SF269 | 12 | 1 | 16.47 | 0 |
| Glycolysis | PTHR21139:SF12 | 10 | 1 | 12.23 | 0 |

**Supplementary Table S28: PANTHER Pathway analysis of PSGs in Asiatic lions**

| PANTHER Pathway | Lion PSGs | PANTHER family/subfamily |
|---|---|---|
| Vitamin B6 metabolism, Threonine biosynthesis | THNSL2 | Threonine synthase-like 2 |
| Nicotine degradation | UGT1A1 | UDP-glucuronosyltransferase 1-1-related |
| Adrenaline and noradrenaline biosynthesis | SLC6A16 | Orphan sodium- and chloride-dependent neurotransmitter transporter NTT5 |
| Ubiquitin proteasome pathway | UBA7 | Ubiquitin-like modifier-activating enzyme 7 |
| PDGF signaling pathway | ARHGAP9 | Rho GTPase-activating protein 9 |
| Ras Pathway | RGL1 | Ral guanine nucleotide dissociation stimulator-like 1 |
| Cadherin signaling pathway | PTPN1 | Tyrosine-protein phosphatase non-receptor type 1 |
| Heme biosynthesis | Coq2 | 4-hydroxybenzoate polyprenyltransferase, mitochondrial |
| Huntington disease | OPTN | Optineurin |
| p53 pathway | Fas | Tumor Necrosis Factor Receptor superfamily member 6 |
| Gq and Go alpha mediated heterotrimeric G-protein signaling pathway | RGS10 | Regulator of g-protein signaling 10 |
| General transcription by RNA polymerase I | TBPL2 | TATA box-binding protein-like protein 2 |
| Apoptosis signaling pathway | Fas, CASP10 | Tumor Necrosis Factor Receptor superfamily member 6, Caspase recruitment domain-containing protein 18-related |
| Inflammation mediated by chemokine | ITGAL, | Integrin alpha-l, |

| | | |
|---|---|---|
| and cytokine signaling pathway | ALOX5AP | Arachidonate 5-lipoxygenase-activating protein |
| Gi and Gs alpha mediated heterotrimeric G-protein signaling pathway | CREB3, RGS10 | Cyclic AMP-responsive element-binding protein 3, Regulator of G-protein signaling 10 |
| FAS signaling pathway | Fas, CASP10 | Tumor Necrosis Factor Receptor superfamily member 6, Caspase recruitment domain-containing protein 18-related |
| Integrin signaling pathway | ITGAL, COL9A2, Itgad | Integrin alpha-L, Collagen alpha-2 (IX) chain, Integrin alpha-D |
| Transcription regulation by bZIP transcription factor | TTF2, TBPL2, CREB3 | Transcription termination factor 2, TATA box-binding protein-like protein 2, Cyclic AMP-responsive element-binding protein 3 |
| T cell activation | Cd3d, CD3E, CD3G | T-cell surface glycoprotein cd3 delta chain, epsilon chain, gamma chain |

**Supplementary Table S29: Rates of heterozygous SNVs among published felid genomes**

| Species | Number of heterozygous SNVs | Rate of heterozygous SNVs | References |
|---|---|---|---|
| **Asiatic lion** | **745,184** | **0.000276** | **This study** |
| African lion | 1,934,590 | 0.000717 | Kim et al. 2016 |
| White lion | 1,630,777 | 0.000604 | “ |
| Bengal tiger | 2,410,975 | 0.000893 | “ |
| Amur tiger | 2,703,974 | 0.001001 | “ |
| White tiger | 2,249,985 | 0.000833 | “ |
| Amur leopard | 1,222,100 | 0.000453 | “ |
| Snow leopard | 1,117,356 | 0.000414 | “ |
| Leopard Cat | 5,625,748 | 0.002084 | “ |
| Cheetah | 1,145,005 | 0.000424 | “ |
| Eurasian lynx | - | 0.000276 | Abascal et al. 2016 |
| Iberian lynx | - | 0.000102 | “ |

**Supplementary Table S30: Male-specific genes* or genes found to be one-fold enriched in three male Asiatic lion transcriptomes.**

| Lion gene | PPDE** | PostFC*** | RealFC**** | UniProt ID | Protein/ Gene name | GO [GO ID] | Protein family |
|---|---|---|---|---|---|---|---|
| **scaffold42437_ size21949.1** | 1 | 175.51 | 13002.75 | K9KBN2 | ATP-dependent RNA helicase DDX3Y-like protein (Fragment) | ATP binding [GO:0005524]; helicase activity [GO:0004386]; nucleic acid binding [GO:0003676] | DEAD box |
| **scaffold59869_ size15355.1** | 1 | 127.61 | 9472.39 | A0A0G2YAU5 | USP9Y | thiol-dependent ubiquitinyl hydrolase activity [GO:0036459]; protein deubiquitination [GO:0016579]; ubiquitin-dependent protein catabolic process [GO:0006511] | Peptidase |
| **scaffold21300_ size35976.1** | 1 | 103.58 | 316.08 | F6RKG6 | ZFX | nucleus [GO:0005634]; metal ion binding [GO:0046872]; RNA polymerase II regulatory region sequence-specific DNA binding [GO:0000977]; transcription factor activity, sequence-specific DNA binding [GO:0003700]; multicellular organism development | NA |

| | | | | | | | |
|---|---|---|---|---|---|---|---|
| | | | | | | [GO:0007275]; regulation of transcription, DNA-templated [GO:0006355] | |
| **scaffold38865_ size23674.1** | 1 | 100.08 | 7381.71 | W8CEP0 | EIF2S3Y | GTP binding [GO:0005525]; GTPase activity [GO:0003924] | NA |
| **scaffold34789_ size25890.1** | 0.99999 | 50.41 | 3686.02 | A0A0G2YEY4 | KDM5D | nucleus [GO:0005634]; DNA binding [GO:0003677]; oxidoreductase activity, acting on paired donors, with incorporation or reduction of molecular oxygen, 2-oxoglutarate as one donor, and incorporation of one atom each of oxygen into both donors [GO:0016706]; zinc ion binding [GO:0008270] | NA |

***Genes with a minimum value of 2 in PostFC,**

****PPDE = posterior probability that a gene/transcript is differentially expressed;**

*****PostFC = posterior fold change for a gene/transcript;**

******RealFC = real fold change for a gene/transcript**
